# A monoclonal antibody targeting the Nipah virus fusion glycoprotein apex imparts protection from disease

**DOI:** 10.1101/2022.09.26.507980

**Authors:** Victoria A. Avanzato, Trenton Bushmaker, Kasopefoluwa Y. Oguntuyo, Kwe Claude Yinda, Helen M. E. Duyvesteyn, Robert Stass, Kimberly Meade-White, Rebecca Rosenke, Tina Thomas, Greg Saturday, Katie J. Doores, Benhur Lee, Thomas A. Bowden, Vincent J. Munster

## Abstract

Nipah virus (NiV) is a highly pathogenic paramyxovirus capable of causing severe respiratory and neurologic disease in humans. Currently, there are no licensed vaccines or therapeutics against NiV, underscoring the urgent need for the development of countermeasures. The NiV surface-displayed glycoproteins, NiV-G and NiV-F, mediate host cell attachment and fusion, respectively, and are heavily targeted by host antibodies. Here, we describe a vaccination-derived neutralizing monoclonal antibody, mAb92, that targets NiV-F. Structural characterization of the fab region bound to NiV-F (NiV-F–Fab92) by cryo-electron microscopy analysis reveals an epitope in the DIII domain at the membrane distal apex of NiV-F, an established site of vulnerability on the NiV surface. Further, prophylactic treatment of hamsters with mAb92 offered complete protection from NiV disease, demonstrating beneficial activity of mAb92 *in vivo*. This work provides support for targeting NiV-F in the development of vaccines and therapeutics against NiV.

## Introduction

The prototypic henipaviruses (HNVs), Hendra (HeV) and Nipah (NiV), are re-emerging, highly pathogenic paramyxoviruses that cause severe neurologic and respiratory disease in humans associated with high morbidity and mortality^1^. NiV has caused near annual outbreaks since 2001 in Southeast Asia^2–5^, with the 2018 NiV outbreak in India exhibiting a 91% case fatality rate^6^. Concerningly, evidence for human-to-human transmission has also been observed ^3,4^. Both viruses circulate in host *Pteropus* bat species and spillover into the human population occurs through intermediate hosts, such as pigs and horses, or directly from bats, primarily through consumption of contaminated date palm sap or fruit^1,4^. Despite these regular NiV and HeV outbreaks with high mortality, there are currently no vaccines or therapeutics against NiV or HeV infection licensed for human use. For these reasons, the WHO has classified NiV as a priority pathogen^7^, underscoring the need for research and development of countermeasures.

The henipaviral surface is decorated with two envelope glycoproteins, the receptor binding protein (HNV-G, also called RBP) and the fusion glycoprotein (HNV-F)^8^. The receptor binding protein is a tetrameric type II integral membrane protein and is responsible for cell attachment through binding to host ephrins, which serve as the functional viral entry receptors^9^. The fusion glycoprotein is a class I fusion protein that is displayed as a trimer on the HNV surface. Similar to other paramyxoviral fusion proteins, crystal structures of NiV-F and HeV-F have demonstrated that HNV-F consists of three domains in the globular head (DI, DII, and DIII), followed by the C-terminal stalk, transmembrane region, and cytoplasmic domain^10–13^. Two canonical heptad repeat (HR) domains are also present. HRA is located in the DIII domain and HRB is located in the stalk region. A third HR domain has recently been described in the N-terminal region of the DIII domain^14^. HNV-F also contains a cleavage site and hydrophobic fusion peptide. Both HeV-F and NiV-F are produced as a full-length precursor, F_0_, which is then endocytosed from the cell surface and cleaved by host-cell cathepsins within the endosomal compartment, forming disulfide linked F_1_ and F_2_ subunits. This process releases the hydrophobic fusion peptide at the N-terminus of the F_1_ subunit, generating fusion-competent HNV-F capable of both virus-cell fusion during entry and cell-cell fusion resulting in syncytia formation^15–17^.

As HNV-G and HNV-F are exposed on the virion surface, these protein antigens are major targets for host-derived antibodies and thus make attractive immunogens^18^. Several promising experimental vaccine candidates utilizing various platforms elicit neutralizing antibodies against the surface glycoproteins^19–24^. Additionally, several *in vivo* studies have demonstrated that passive transfer of vaccine serum or administration of monoclonal antibodies targeting the attachment or fusion glycoproteins, both prophylactically and post-infection, protects against henipaviral disease in hamster, ferret, and African green monkey models^25–32^. Hybridoma-secreted mAbs targeting HNV-F have been shown to protect hamsters from disease at low doses^25^. One humanized HNV-F targeting antibody, mAb 5B3, protects ferrets from NiV and HeV disease when administered several days after challenge^30^. Additionally, the HNV-G-targeting monoclonal antibody mAb 102.4 protects against NiV and HeV disease in the African green monkey model^28,29^ and has recently been shown to be safe in a Phase I clinical trial in Australia^33,34^, highlighting the potential clinical utility of such interventions.

The promise demonstrated by antibody mediated countermeasures has prompted us, as well as others, to study epitopes on the henipaviral glycoproteins in more detail^31,32, 35–39^. Previously, we solved the crystal structure of mAb66 bound to the prefusion NiV-F trimer, which revealed a site of vulnerability at the fusion protein apex within the DIII domain^35^. Additionally, we observed a high level of sequence conservation at the epitope as well as the presence of structural and functional constraints, supporting that this region makes an attractive target for vaccination-derived or therapeutic antibodies. To further expand the understanding of antibody-mediated targeting of HNVs and identify additional epitopes on the fusion glycoprotein, we determined the structure of an immunization-derived neutralizing monoclonal antibody that targets HNV-F, termed mAb92. Our structure reveals that mAb92 recognizes a similar but distinct epitope to that of mAb66 at the NiV-F apex. Additionally, we show that prophylactic treatment with mAb92 protects Syrian hamsters from a lethal NiV challenge, further supporting the potential utility in further development of vaccines and therapeutics targeting HNV-F.

## Results

### Structural characterization of Fab92 in complex with NiV-F by cryo-electron microscopy

mAb92 is a neutralizing monoclonal antibody that targets NiV-F and was derived through DNA vaccination of rabbits with expression plasmids encoding codon-optimized NiV-F and wild-type NiV-G and NiV-M^40^. mAb92 neutralizes NiV-F/G mediated viral entry and fusion with an IC_50_ value of <1 μg/mL^40^. For structural analysis, mAb92 was purified from hybridoma supernatant by Protein G purification. The Fab92 region was isolated via papain cleavage and purified by size exclusion chromatography (SEC), then incubated in excess with recombinantly produced NiV-F to allow for complexation (Fig S1). The NiV-F–Fab92 complex was purified for structural characterization by single particle cryo-EM (Materials and Methods).

Single particle cryo-EM analysis and refinement generated a reconstruction of the NiV-F–Fab92 complex with a reported global resolution of 3.5 Å (GSFSC=0.143, cryoSPARC v3.2.0^41,42^, Fig S2). In our reconstruction, we observed a single Fab92 molecule bound to the NiV-F trimer (Fig 1A). The local resolution was highest for the Fab92 variable regions and the NiV-F trimer (Fig S2). The Fab92 constant domains were poorly resolved, which precluded their modelling. The sequence of the Fab92 variable region was rescued from hybridomas to allow building of the variable regions (Fig S3). The final model thus included the Fab92 variable regions and the NiV-F ectodomain residues I27 – A477, with a chain break at the cleavage site/fusion peptide, residues V105 – A111, and a loop containing E163 – E166, as these residues were not resolved. The structure reveals a single Fab92 molecule bound to an epitope at the apex of the membrane distal DIII domain of the NiV-F trimer (Fig 1). Data collection and refinement statistics are shown in Supplementary Table S1 (SI Table S1).

**Figure 1.**
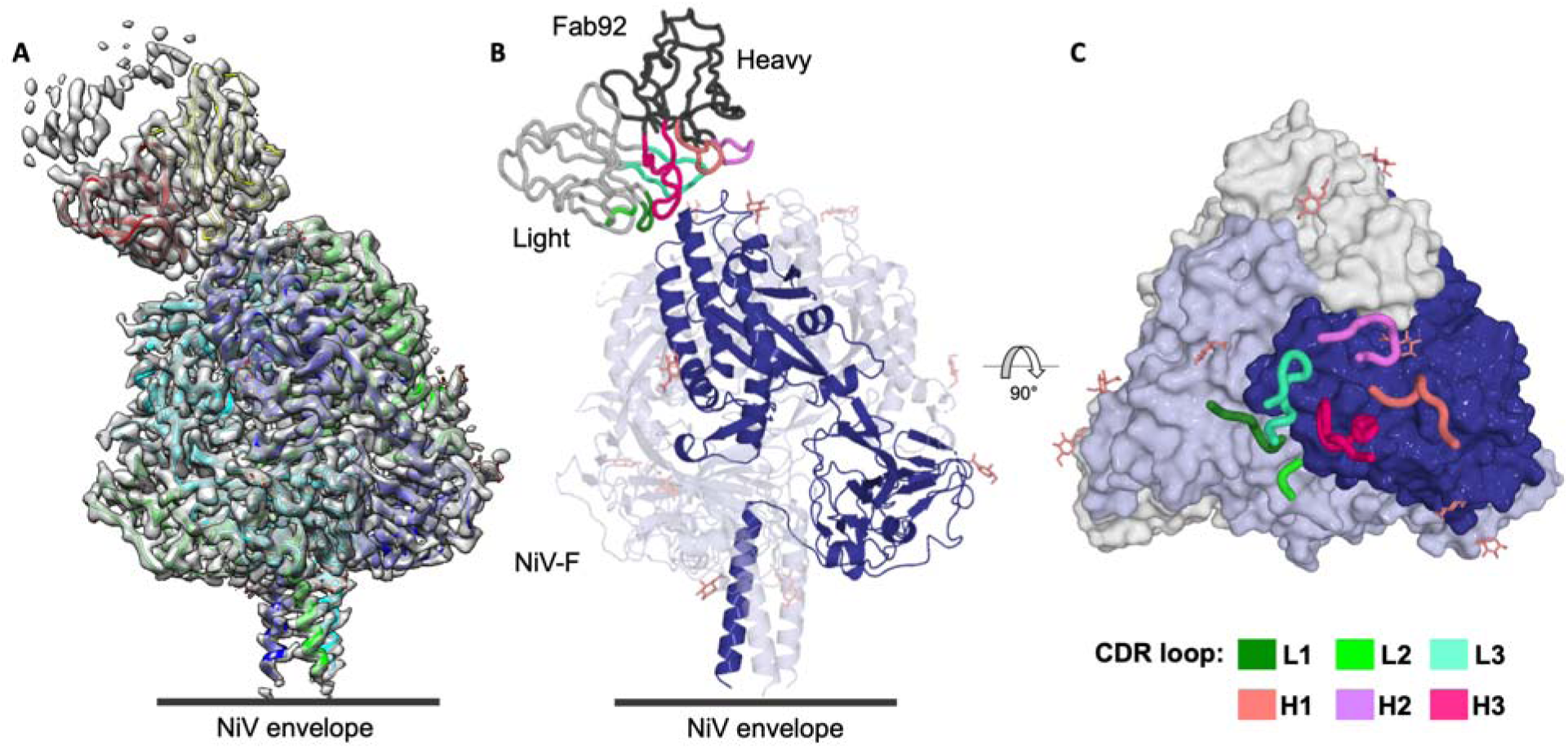
The cryo-EM structure of the NiV-F–Fab92 complex. (A) The model of the NiV-F– Fab92 complex fit into the cryo-EM map (transparent surface). The relative position of the NiV envelope is shown as a black line. (B) Side view of the NiV-F–Fab92 complex, shown in cartoon representation. The NiV-F trimer is colored light blue, with the F protomer to which Fab92 is bound colored dark blue for clarity. The Fab92 heavy chain is colored dark gray and the light chain is colored light gray. Heavy chain CDRs are colored pink and light chain CDRs are colored green, according to the legend. (C) Top view of the NiV-F–Fab92 complex. The NiV-F trimer is shown with surface representation and each F protomer is colored a different shade of blue. The F protomer to which Fab92 is bound is dark blue. The Fab92 CDR loops are shown as tubes, and the heavy and light CDRs are colored shades of pink and green according to the legend. Glycans observed in the reconstruction are depicted as sticks.

### mAb92 targets an epitope at the apex of NiV-F

Both the Fab92 heavy and light chains contribute to a roughly ∼600 Å^2^ interface at the apex of NiV-F. The compact epitope primarily comprises the region above the central helix on a single protomer of the NiV-F trimer and includes residues within the ranges of Asn67 – Ser74 and Ile187 – Glu196, (Fig 1B and C). Individually, CDR H3 produces the largest interface of all the Fab92 CDRs by interacting with residues ranging between Ser69 – Cys71 and Ile187 – Cys192 on NiV-F (Fig 2A, Fig S4). The backbone of CDR H3 residue Pro97 and Gly99 both form hydrogen bonds with the side chain and backbone of NiV-F residue Ser69. CDR H3 residues Pro97 and Gly100A (Chothia numbering scheme^43^) also form hydrogen bonds with residue Gln70 on NiV-F. Additionally, the sidechain of H3 Tyr100B is occluded within the antibody-antigen interface, forming stabilizing H-bonds with NiV-F Ile187 and Cys71. Residue Tyr33 on CDR H1 also forms contacts with NiV-F Gln70. While CDR H2 does not form appreciable protein contacts with NiV-F, residue Tyr58 in heavy chain framework regions 3 (FR3), immediately adjacent to CDR H2, contacts residue Asp188 on NiV-F (Fig 2A).

**Figure 2.**
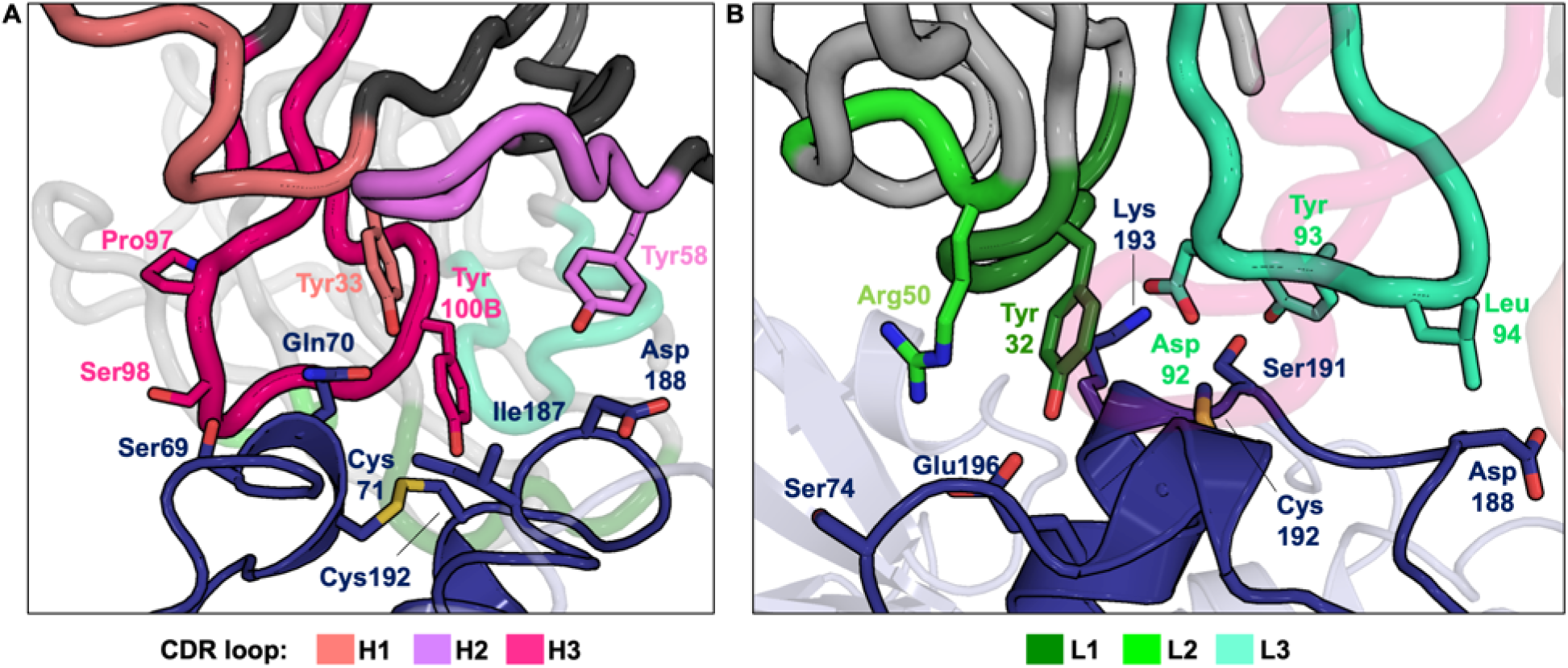
Molecular characterization of the Fab92 epitope. (A) Close up view of the heavy chain CDR molecular interactions with NiV-F. (B) Close up view of light chain CDR molecular interactions with NiV-F. Fab92 is shown as a gray cartoon tube and NiV-F is shown as a dark blue cartoon. F protomers not bound by Fab92 are shown as light blue and slightly transparent. CDR loops are colored in shades of green (light chain) and pink (heavy chain), as indicated. Sidechains of residues participating in interactions such as intermolecular hydrogen bonds or salt bridges are shown as sticks.

Fab92 light chain CDR L3 residues Asp92, Tyr93, and Leu94 extensively contact Ile187 – Glu196 on NiV-F. In particular, L3 Asp92 forms both backbone and sidechain H-bond interactions with residues Cys192 and Lys193 on NiV-F, and an additional salt bridge with the sidechain of NiV-F Lys193. On CDR L2, Arg50 forms an H-bond and salt bridge with the side chain of Glu196 on NiV-F. Additionally, the sidechain from residue Tyr33 on CDR L1 forms stabilizing H-bond interactions with NiV-F Cys192 and Glu196, and contacts NiV-F Lys193 (Fig 2B).

### Glycosylation at the epitope on NiV-F affects mAb92 binding

Of the five encoded putative N-linked glycosylation sites on NiV-F (Asn64, Asn67, Asn99, Asn414, and Asn464, named F1 – F5, respectively), four have been shown to be occupied on recombinant soluble protein^13,35,36^ as well as on NiV-F present on infectious pseudotyped particles^44,45^. Consistent with these experimental observations, there was evidence for glycosylation at four of the five potential glycosylation sites (F2 – F5) in the cryo-EM density of the NiV-F structure. There was no density observed at the F1 glycan site. Due to the inherent flexibility expected of glycan chains, the observable density did not permit modelling beyond the first GlcNAc residue at each of the F2-F5 glycan sites on NiV-F.

Similar to previous structural observations with another anti-NiV-F neutralizing mAb, mAb66^35^, the F2 glycan is present at the periphery of the Fab92 epitope. Both the Fab92 heavy and light chains contact the glycan at the F2 site on both the corresponding and neighboring protomer to which Fab92 is bound (Fig 3B). While CDR H2 does not contact proteinaceous residues at the NiV-F epitope, H2 residue Thr53 contacts the F2 glycan on the corresponding NiV-F protomer. On the light chain, residues Gln27 from CDR L1 and Tyr93 from CDR L3 contact the F2 glycan on the neighboring protomer (Fig 3B). Modelling a full-length complex-type glycan on to this site shows the glycan chain extending outwards from the Fab92 heavy and light chains, allowing Fab92 access to its proteinaceous epitope on the F protomer (Fig 3C).

**Figure 3.**
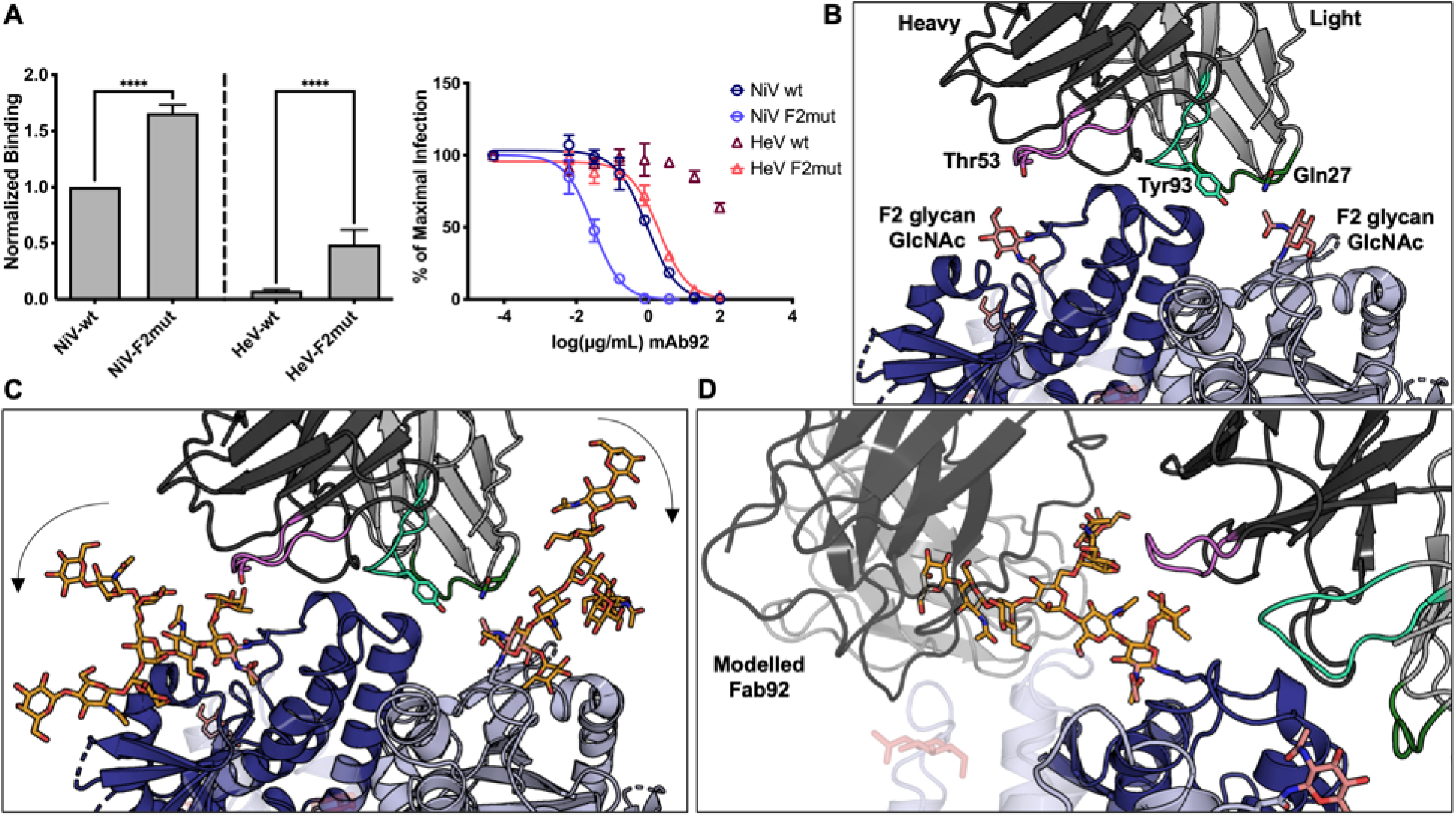
Glycosylation at the F2 site on NiV-F impedes Fab92 binding. (A, left panel) mAb92 binding of NiV-F or HeV-F in the presence or absence of the F2 glycan (WT and F2 mut, respectively). 293T cells were transiently transfected to express the indicated HNV-F glycoprotein prior to flow cytometry with mAb92. Binding was normalized to 2 anti-HNV-F polyclonal antibodies, pAb2489 and pAb2490, previously shown to have equivalent cross reactivity to NiV and HeV-F WT and mutant constructs^35^, as indicated in the Methods. The polyclonal antibodies were further normalized to WT NiV-F binding for further comparison. Presented are the results from three independent biological replicates with statistics determine by one-way ANOVA with Sidak’s correction for multiple comparisons (****p< 0.0001). (A, Right panel) mAb92 neutralization of HNV pseudotyped particles (HNVpp) infection on U87 MG glioblastoma cells HNVpp bearing WT-G and WT-F (F2 glycan present) or F2mut (F2 glycan absent) glycoproteins were produced with a VSVΔG-RLuc system as described in the Methods. HNVpp input was optimized to give output within the dynamic range of the assay. A constant amount of the indicated HNVpp was used to infect permissive U87 cells in the presence or absence of a serial 5-fold dilution of mAb92. Data presented are from three independent biological replicates, each with technical duplicates, and points are presented as the mean +/- SE. Neutralization curves were constrained by artificially setting the lowest mAb concentration (X-axis) to maximum infection (Y-axis) to represent neutralization in the absence of any mAb. The data were then analyzed using a variable slope model with a 4-parameter dose-response curve (GraphPad PRISM), and statistical significance was tested with a 2-way ANOVA with Dunnett’s correction for multiple comparisons. (B) Models for Fab92 and NiV-F are shown in cartoon representation and colored gray and blue, respectively. The NiV-F protomer to which Fab92 is bound is colored dark blue and the neighboring protomers are colored light blue. Residues from CDR loops L1 (dark green), L3 (light green), and H2 (pink) contact the first GlcNAc residue of the F2 glycan on two protomers of NiV-F. The sidechains of residues observed to interact with the first GlcNAc of the F2 glycan (as calculated by the PDBePISA server^46^) are shown as sticks and colored according to CDR loop as described in Figure 1. The sidechain of the F2 Asn67 residue is shown as blue sticks. The first F2 GlcNAc is shown as salmon sticks. (C) The complex glycan (shown as light orange sticks) from the structure PDB 4BYH^65^ was modelled onto the F2 glycan site on two protomers of NiV-F by alignment onto the first GlcNAc that was observed in the cryo-EM-derived reconstruction. The black arrows indicate the direction of the F2 glycan extending outwards away from Fab92, allowing Fab92 to access its proteinaceous epitope. (D) To investigate if the presence of a full-length F2 glycan may interfere with Fab92 binding to adjacent protomers, Fab92 was superposed onto the binding site on the adjacent NiV-F protomers. The Fab92 modelled onto the adjacent protomer is colored gray. The modelled glycan at the F2 position is shown to clash with a modelled Fab92 bound to the adjacent protomer.

To further investigate the effect of the F2 glycan on mAb92 binding, Fab92 was superposed onto the unoccupied NiV-F protomer binding sites of the F2 glycosylated NiV-F trimer. This analysis revealed that the modeled glycan at the F2 site extends into the Fab92 binding site on the neighboring protomer and clashes with the modelled Fab92 (Fig 3D). Thus, it is possible that binding of one Fab92 molecule to the NiV-F trimer repositions the F2 glycan such that it obstructs the adjacent Fab92 binding sites of the other NiV-F protomers and may explain the observation of a single Fab92 molecule in the 2D classes and reconstruction (Fig 1, Fig S2).

Consistent with the structural observations, the removal of the F2 glycan site on NiV-F and HeV-F (NiV-F2mut and HeV-F2mut) significantly enhances binding by mAb92 (Fig 3A, left), as well as neutralization of pseudoparticles (Fig 3A, right). Notably, mAb92 is capable of binding HeV-F and neutralizing HeV pseudoparticles in the absence of the F2 glycan, suggesting a role of this glycan in epitope shielding. These findings are similar with the observations that another reported HNV-F targeting mAb, mAb 12B2, also displayed increased binding in the absence of the F2 glycan^37^. This contrasts previous observations that mAb66 binding to HNV-F was not observably affected by the removal of the F2 glycan^35^, though this did increase mAb66 mediated neutralization of pseudoparticles, revealing the differential dependence of monoclonal antibodies upon NiV-F glycosylation.

### Difference between NiV and HeV at the mAb92 epitope

Previous functional studies have shown that mAb92 binds more strongly to NiV-F than HeV-F^40^. Mapping of the Fab92 epitope onto an amino acid sequence alignment of the NiV and HeV fusion proteins reveals three residues that differ within the epitope between NiV-F and HeV-F. Residues Gln70, Ser74, and Lys189 in NiV-F are replaced by Lys70, Thr74, and Gln189 in HeV-F, respectively (Fig 4A and B). While Ser74 and Lys189 on NiV-F make relatively minor contributions to the Fab92 epitope (as calculated by the PDBePISA server^46^), residue Gln70 on NiV-F makes substantial contacts with Fab92. In particular, residues Pro97 and Gly100A in Fab92 CDR H3 form hydrogen bonds with the Gln70 sidechain and backbone. Tyr33 from CDR H1 also forms extensive contacts with Gln70 (Fig 4C). Modelling of Fab92 bound to HeV-F reveals that certain lysine rotamers generated by the Gln to Lys substitution at amino acid position 70 introduce steric clashes with Fab92 residues Tyr33 and Tyr100B on CDR H1 and H3. Such clashes would likely hinder the ability of Fab92 to bind HeV-F (Fig 4D), as observed for mAb66 previously^35^.

**Figure 4.**
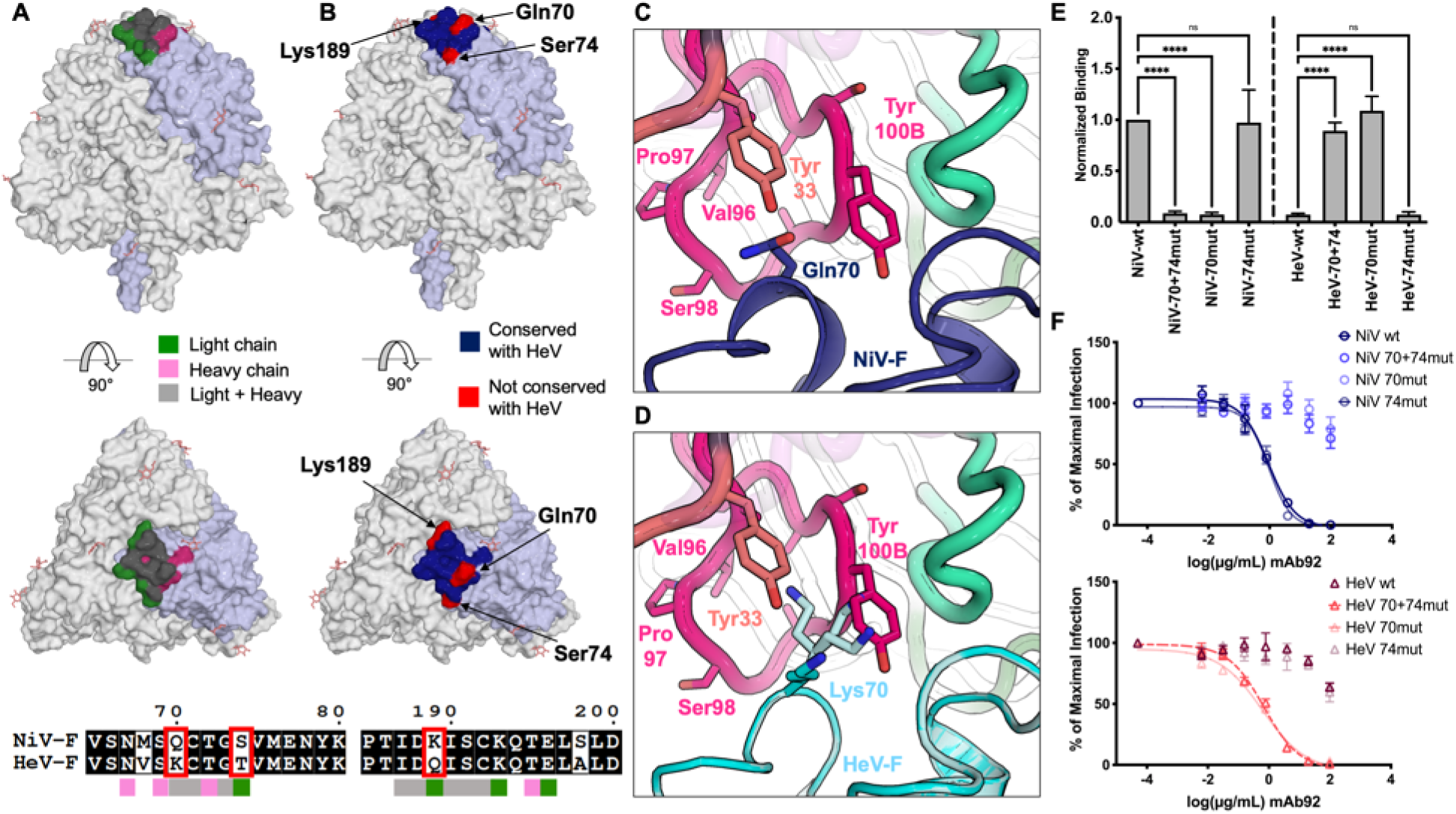
Differences between NiV-F and HeV-F within the Fab92 binding footprint. (A) The residues on the NiV-F surface in the binding interface are colored according to the Fab92 chain involved in the contact. Residues contacted by the light chain are highlighted green, residues contacted by the heavy chain are highlighted pink, and those contacted by both chains are shown in gray. The NiV-F trimer is depicted with surface representation in white, with a single protomer that contains the Fab92 epitope in light blue for clarity. (Lower) Amino acid sequence alignment between NiV-F (Malaysia, AAV80428.1) and HeV-F (AEB21197.1) was generated by Multalin^66^ and plotted by ESPript^67^. Residues involved in the Fab92 interface are annotated with colored boxes below the alignment. Residues in the interface that differ between NiV-F and HeV-F are outlined by a red box. (B) Surface view of the NiV-F trimer shown in white, with the protomer with Fab92 bound shown in light blue. Residues contacted by Fab92 are colored dark blue. Residues in the epitope that are conserved between NiV-F and HeV-F are colored dark blue and those that differ are colored red. (C) Molecular interactions of Fab92 with NiV-F residue Gln70. Sidechains of CDR H3 (hot pink) and Gln70 are shown as sticks. NiV-F is shown in dark blue. (D) A model for steric hinderance in Fab92 recognition of HeV-F imposed by Lys70. To better understand the interactions between Fab92 and HeV-F, the HeV-F structure (PDB ID 5EJB^12^, cyan cartoon) was aligned to NiV-F using COOT. The lysine at position 70 is modelled as observed in the HeV-F crystal structure, as well as in possible rotamer configurations that introduce steric clashes with Fab92 residues Tyr33 (CDR H1, pink) and Tyr100B (CDR H3, hot pink). (E) mAb92 binding of NiV and HeV-F WT or mutant F proteins. 293T cells were transfected as described in the Methods to express WT HNV-F, the double residue 70 and 74 mutant (70+74mut), the residue 70 only mutant (70mut), or the residue 74 only mutant (74mut). Data presented are the result of three independent biological replicates and are analyzed as described previously for Fig 3A, left panel (ns, not significant, **** p < 0.0001). (F) mAb92 neutralization of HNVpp expressing WT or mutant F glycoproteins. HNVpp were produced as described in the Methods to express WT G and the indicated NiV/HeV WT or mutant (70+74mut, 70mut, or 74mut) F glycoprotein. Neutralization curves were generated from three independent biological replicates done in technical duplicates. Data were analyzed as described above for Fig 3A, right panel. WT data for binding and neutralization are repeated from Fig 3A to facilitate interpretation of these findings in the context of the 70 and 74 double or individual mutants.

To further evaluate the functional effects of these substitutions on mAb92 binding of HNV-F and neutralization of HNVs, reciprocal mutants were generated, in which residues Gln70 and Ser74 of NiV-F were replaced with Lys70 of and Thr74 HeV-F, both together and individually. The cell based binding assay confirmed that substitution of the mutations Gln70 and Ser74 in NiV-F to Lys70 and Thr74 in HeV-F (Q70K and S74T, NiV-70+74mut) significantly impairs binding of mAb92 (Fig 4E). Consistent with the structural predictions, mAb92 binding was significantly impaired when NiV-F Gln70 was replaced with Lys70 (NiV-70mut), whereas substitution of NiV-F Ser74 with HeV Thr74 (NiV-74mut) did not affect mAb92 binding. The converse was also true, as a single point mutation in HeV-F, replacing Lys70 with Gln70 (HeV-70mut), restored mAb92 binding to levels nearly comparable with that of NiV-wt (Fig 4E). These findings are also corroborated in the pseudotyped particle neutralization assay (Fig 4F), highlighting the importance of residue Gln70, but not Ser74, for binding and neutralization by mAb92. Interestingly, Dang et al^37^ postulate that a K70Q mutation in HeV-F confers enhanced binding by mAb 12B2, suggesting mAbs 92 and 12B2 share a reliance on NiV-F residue Gln70 for binding and neutralization.

### Prophylactic treatment with mAb92 protect Syrian hamsters from lethal Nipah virus challenge

To determine whether mAb92 displayed therapeutic activity *in vivo*, prophylactic treatment of mAb92 was evaluated in the Syrian hamster model. Four groups of 11 animals were included in the experiment; groups I and III received mAb92 and groups II and IV received saline. Groups I and II were then challenged with NiV, and groups III and IV with HeV, and monitored for antibody titers, viral titers, clinical disease, and survival.

No pre-existing antibodies cross-reactive to NiV-F were detectable prior to the start of the study in any animal (Fig 5A). Two days before lethal challenge with NiV or HeV (-2 DPI, Days Post Inoculation), hamsters in groups I and III received mAb92 (500 μg per hamster) and those in groups II and IV received saline as controls, via the intramuscular (I.M.) route. At -1 DPI, hamsters receiving mAb92 displayed NiV-F ELISA titers ranging from 1/12,800 to 1/51,000, suggesting high levels of serum antibody at the time of virus challenge at 0 DPI (Fig 5A). All hamsters then received a lethal challenge dose (100 LD_50_) of either NiV Malaysia (Groups I and II, 6.8 x 10^3^ TCID_50_) or HeV (Groups III and IV, 6.0 x 10^2^ TCID_50_) intraperitoneally (I.P.) at 0 DPI. The prophylactic mAb92 treatment fully protected hamsters from lethal NiV Malaysia challenge and statistical analysis showed that survival in the mAb92 treated group was significant compared to controls (p < 0.01). All seven hamsters in the mAb92 treatment group (Group I) survived until the end of the experiment (28 DPI) and did not show any clinical signs of disease following NiV infection (Fig 5B and Fig S5). Five of the seven control animals (Group II) developed clinical signs of NiV disease, including weight loss, ruffled fur, and hunched posture, and met euthanasia criteria within the first ten days following challenge (Fig 5B and Fig S5). In contrast, mAb92 did not protect the hamsters against lethal HeV challenge, and no significant difference in survival was observed between mAb92 treated (Group III) and control animals (Group IV) following HeV infection, with two treated and one control animal surviving the challenge (Fig 5B). Both HeV groups showed clinical signs of disease, including weight loss, ruffled fur, and hunched posture in the first ten days following challenge (Fig S5).

**Figure 5.**
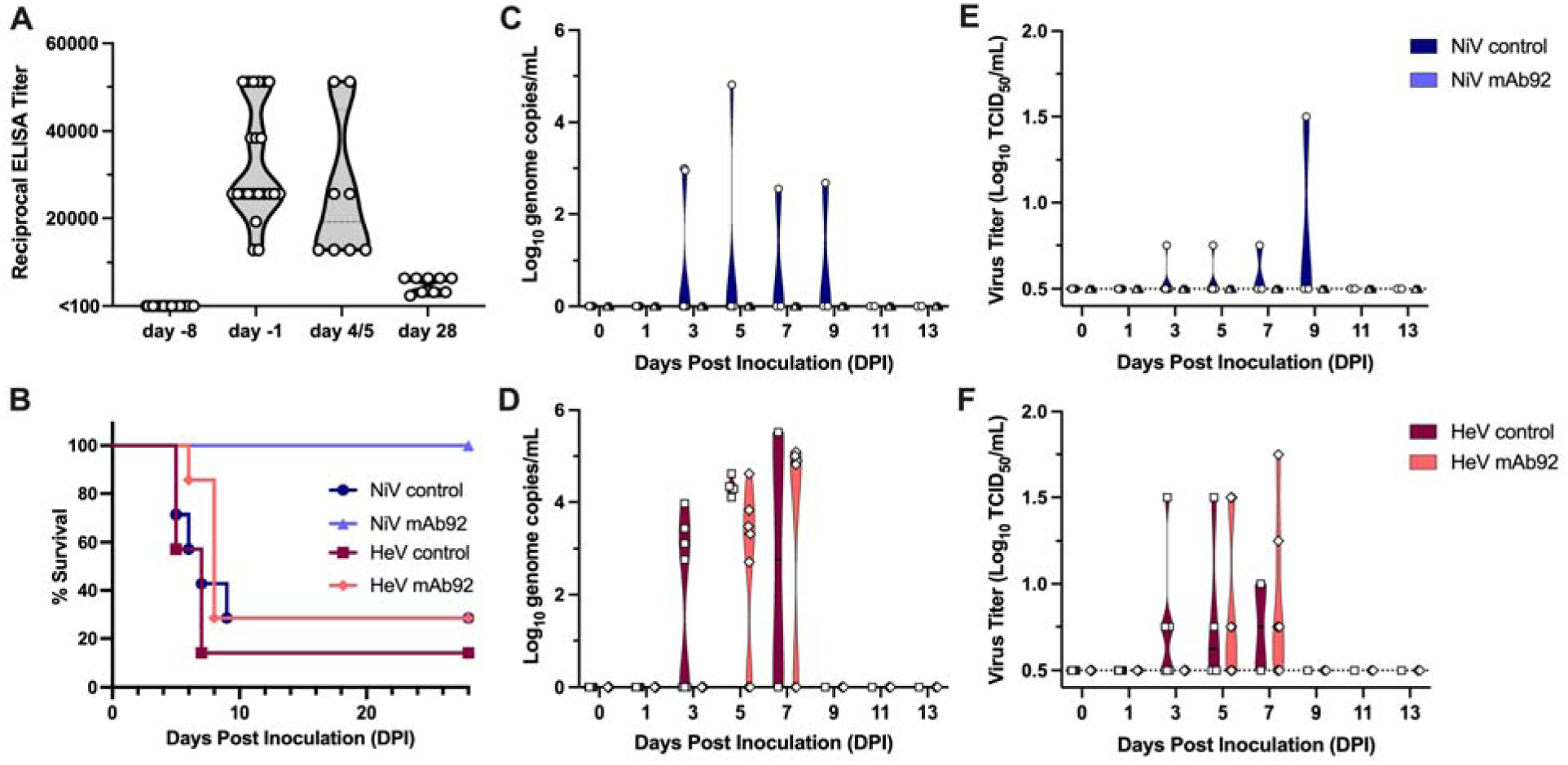
Prophylactic treatment with mAb92 protects Syrian golden hamsters from NiV but not HeV disease. (A) mAb92 titers were determined by ELISA assay against purified NiV-F using anti-rabbit secondary antibody. Each sample was tested in duplicate and each data point represents the end-point titer for an individual hamster. (B) Survival of control hamsters and mAb92 treated hamsters following NiV Malaysia and HeV challenge. Statistical significance was calculated with a Log-rank (Mantel-Cox) test. Each group consisted of seven animals monitored for survival. (C) Viral load (genome copies/mL) in oropharyngeal swabs following challenge with NiV Malaysia determined by qRT-PCR. (D) Viral load (genome copies/mL) in oropharyngeal swabs following challenge with HeV determined by qRT-PCR. (E) Infectious virus titer (TCID_50_/mL) in oropharyngeal swabs collected following challenge with NiV Malaysia determined by end-point titration on Vero E6 cells. (F) Infectious virus titer (TCID_50_/mL) in oropharyngeal swabs collected following challenge with HeV determined by end-point titration on Vero E6 cells. For E and F, the dotted line represents the limit of detection for the titration assay and each point represents a swab from an individual animal at the given time point. In each plot, NiV control animals are represented by dark blue and circles, NiV mAb92 animals by light blue and triangles, HeV control animals by maroon and squares, and HeV mAb92 animals by salmon pink and diamonds.

To assess viral shedding, combined nasal-oropharyngeal swabs were taken every other day until 13 DPI and assessed for the presence of viral RNA (Fig 5C and 5D) and infectious virus (Fig 5E and 5F). Of the animals challenged with NiV, no shedding was detected in any of the mAb92 treated animals throughout the experiment, while swabs from control animals were positive for both NiV RNA and infectious virus from 3 DPI to 9 DPI (Fig 5C and 5E). Within the groups challenged with HeV, control animals began shedding by 3 DPI and the mAb92 treated animals began shedding at 5 DPI, as indicated by detection of both HeV RNA and infectious HeV in the swabs (Fig 5D and 5F). There was no evidence of shedding beyond 9 DPI in any of the surviving animals. This indicates that prophylactic mAb92 treatment prevented viral shedding following challenge with NiV, but not HeV.

Four animals from each group were euthanized at 4 DPI (HeV) or 5 DPI (NiV) and lung and brain tissue were collected for virological and pathological evaluation. In the NiV challenged animals, the group receiving mAb92 displayed significantly lower levels of NiV RNA in both the brain and lung, and no detectable infectious virus in either tissue (Fig 6A and 6B). Following HeV challenge, mAb92 treated animals trended toward lower levels of HeV RNA in the lung and brain, however the difference was not significant (Fig 6C). No infectious virus was detected in the brain of the mAb92 animals, and only one of the mAb92 animals had detectable infectious HeV in the lung (Fig 6D). The lack of significance in the difference in infectious titers in brain tissue in both NiV and HeV groups could be due to inconsistent viral dissemination to the central nervous system in the model using this virus dose; two control animals had positive brain titers for each NiV and HeV. These results show that the prophylactic mAb92 treatment was effective in significantly reducing viral RNA in lung and brain tissue following NiV challenge, and infectious viral load in lung tissue following infection with both NiV and HeV. The reduction is more robust in the NiV group indicating that mAb92 is more effective against NiV than HeV, consistent with previous *in vitro*^40^ and structural analyses (Fig 4). The mAb92 titers in these animals remained elevated at 4 and 5 DPI, with titers again ranging from 1/12,800 to 1/51,200 (Fig 5A).

**Figure 6.**
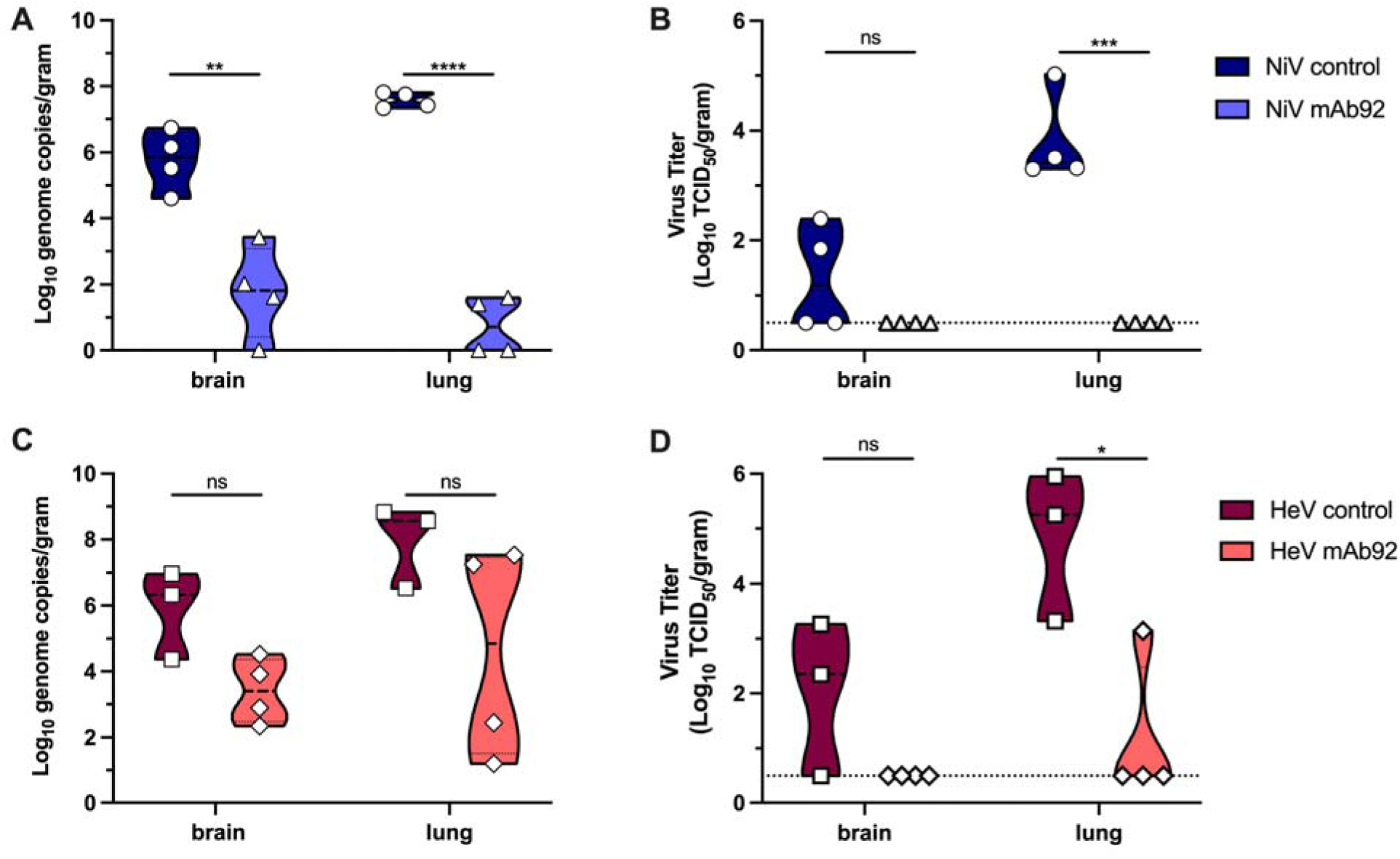
Prophylactic treatment with mAb92 reduces viral loads in tissues from hamsters following challenge with NiV and HeV. (A) Viral load (genome copies/gram) in lung and brain tissue collected from hamsters challenged with NiV Malaysia at 5 DPI determined by qRT-PCR. (B) Infectious virus titer (TCID_50_/gram) in lung and brain tissue collected from hamsters following NiV Malaysia challenge at 5 DPI, determined by end-point titration on Vero E6 cells. (C) Viral load (genome copies/gram) in lung and brain tissue collected from hamsters challenged with HeV at 4 DPI, determined by qRT-PCR. (D) Infectious virus titer (TCID_50_/gram) in lung and brain tissue collected from hamsters following HeV challenge at 4 DPI, determined by end-point titration on Vero E6 cells. For C and D, a dotted line represents the limit of detection for the titration assay. Statistical significance was determined using an unpaired t-test in GraphPad Prism, v.9.1.0. Significance is denoted as follows: ns = p > 0.05, * = p ≤ 0.05; ** = p ≤ 0.01; *** = p ≤ 0.001; **** = p ≤ 0.0001. In each plot, NiV control animals are represented by dark blue and circles, NiV mAb92 animals by light blue and triangles, HeV control animals by maroon and squares, and HeV mAb92 animals by salmon pink and diamonds.

Lung tissue collected at 4 DPI (HeV) and 5 DPI (NiV) was then evaluated for gross pathological and histological changes (Fig 7). Control animals from both NiV and HeV groups displayed gross pathological lesions including red, edematous lungs. The mAb92 treated animals in the NiV groups did not display any gross pathological lesions and appeared healthy (Fig 7A). In the HeV group, half of the mAb92 animals displayed lung lesions, whereas the other half did not display any gross pathological lesions and appeared healthy (Fig 7B). Lung tissues were also sectioned and embedded for staining with hematoxylin and eosin (H&E stain, Fig 7C and 7D) and analysis by *in situ hybridization* (ISH, Fig 7E and 7F). The H&E staining revealed animals in both NiV and HeV control groups developed interstitial pneumonia characterized by multifocal inflammatory nodules that are centered on terminal bronchioles with extension into adjacent alveoli. The nodules are composed of large numbers of foamy macrophages and fewer neutrophils and lymphocytes admixed with small amounts of necrotic debris. In most cases there is hemorrhage, fibrin and edema often extend into surrounding alveoli. By contrast, the mAb92 animals in the NiV group showed no signs of pulmonary pathology, while in the HeV group, two of the animals showed no pathological changes and two animals resembled controls. Viral RNA was detected by ISH in all control animals, predominantly in type I pneumocytes but also in vascular endothelium. Consistent with the H&E, no viral RNA was detected in animals treated with mAb92 in the NiV group, while HeV RNA could be detected in the lungs of two mAb92 animals. Together, these observations show that mAb92 may delay the onset of HeV infection, but ultimately did not protect the hamsters from severe disease and death, whereas hamsters were fully protected from the NiV challenge.

**Figure 7.**
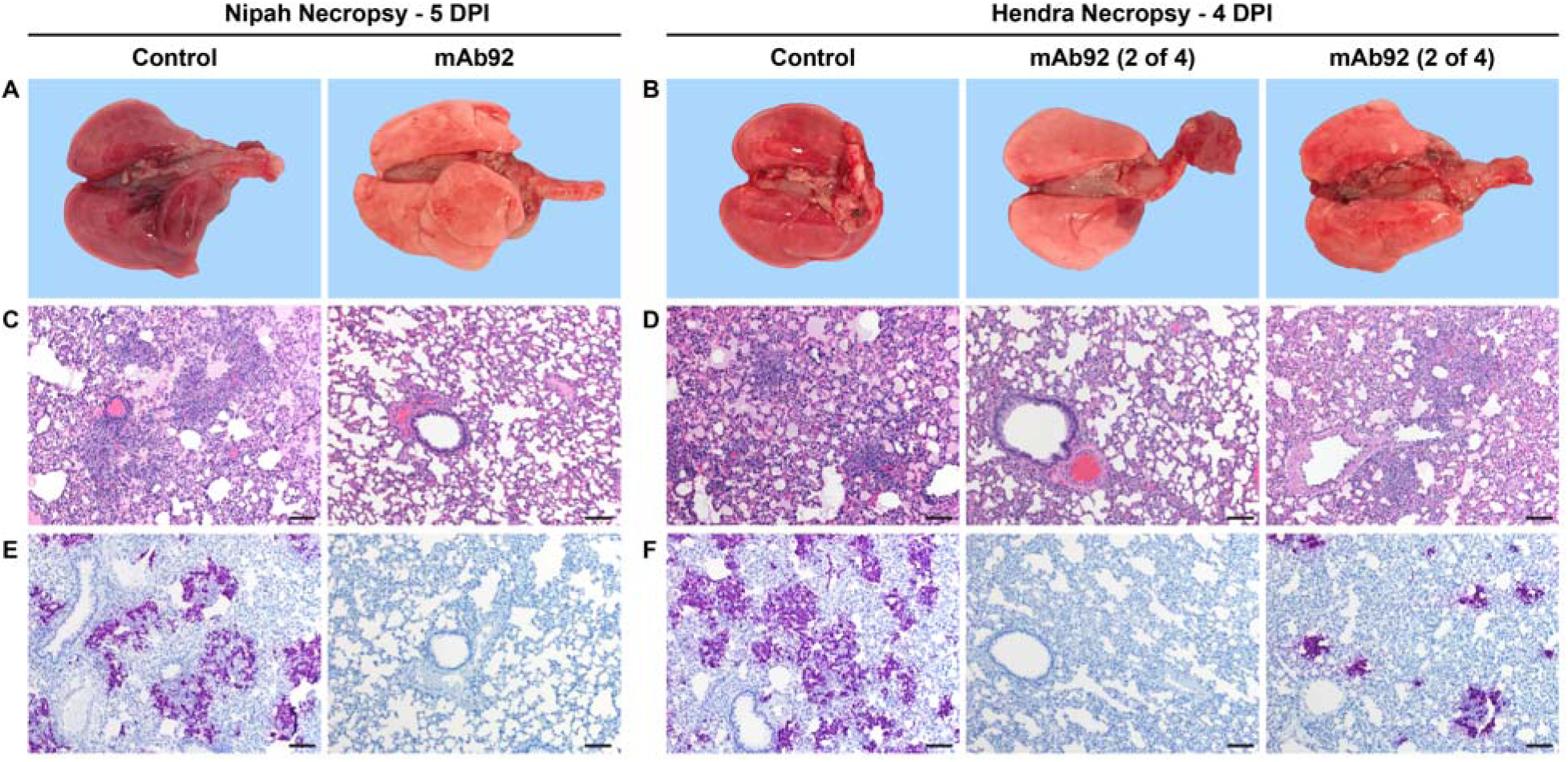
Pathological and histological changes in lung tissue in control and mAb92 treated hamsters after NiV Malaysia or HeV challenge. (A) Gross pathology of lungs from representative control animals (left) and mAb92 treated animals (right), following challenge with NiV Malaysia, euthanized at 5 DPI. (B) Gross pathology of lungs from representative control animals (left) and mAb92 treated animals (middle and right), following challenge with HeV, euthanized at 4 DPI. (C) Hematoxylin and eosin (H&E) staining of lung tissue from representative control animals (left) and mAb92 treated animals (right), following challenge with NiV Malaysia, euthanized at 5 DPI (100x bar = 100 μm). (D) H&E staining of lung tissue from representative control animals (left) and mAb92 treated animals (middle and right), following challenge with HeV, euthanized at 4 DPI (100x bar = 100 μm). (E) *In situ* hybridization (ISH) stain of representative lung from control animals (left) and mAb92 treated animals (right), following challenge with NiV Malaysia, euthanized at 5 DPI (100x bar = 100 μm). (F) ISH stain of representative lung from control animals (left) and mAb92 treated animals (middle and right), following challenge with HeV, euthanized at 4 DPI (100x bar = 100 μm).

Additionally, at 28 DPI, serum from the surviving hamsters in the NiV group were assessed for hamster antibodies against the NiV-G, F, and N proteins to evaluate if the hamsters seroconverted, indicating the generation of an immune response following viral replication. The two surviving control animals developed high titers against all three NiV proteins. In contrast, the mAb92 treated animals had low to no detectable titers against NiV-F, G, and N, which suggests very little, if any, NiV replication occurred following challenge in the mAb92 treated group (SI Table S2). Detectable mAb92 titers were substantially decreased at 28 DPI, ranging from 1/2400 to 1/6400 (Fig 5A).

## Discussion

NiV continues to re-emerge, as recently as 2021 in Kerala India^47^, causing highly fatal outbreaks and underscoring the need for research to develop countermeasures. Several *in vivo* studies have demonstrated that neutralizing antibodies against the NiV surface glycoproteins offer protection from disease^25–30^. Structural characterization of several monoclonal antibodies targeting HNV-G^31,32,38^ and HNV-F^35–37,39^ has allowed detailed descriptions of the molecular basis by which NiV can be neutralized via antibody targeting of the surface glycoproteins. Here, we provide a structural and functional characterization of mAb92, a neutralizing antibody that targets NiV-F generated through the vaccination of rabbits, and show that prophylactic mAb92 treatment protects hamsters from lethal NiV disease.

Structural characterization of mAb92 by cryo-electron microscopy reveals an epitope at the apex of the DIII domain on the NiV-F trimer, similar to the previously described epitopes of mAb66^35^ and 12B2^37^. The apical residues contained in these described epitopes are highly conserved amongst NiV Malaysia and Bangladesh strains. However, three residues differ in the mAb92 epitope between NiV and HeV, which likely contribute to the decreased binding and neutralization potency of mAb92 for HeV. In particular, the Gln to Lys substitution at position 70 likely contributes to this observation, as mAb92 forms extensive interactions with residue Gln70 on the NiV surface, similar to previous observations with mAb66^35^. This apical region has been shown to be a common target on HNV-F and displays high genetic conservation suggestive of functional constraints, supporting this region as a site of vulnerability and an attractive target for development of vaccines and therapeutics^35^. Notably, most anti-HNV-F mAbs characterized to date target membrane-distal epitopes within DIII (currently available structures are summarized in Fig 8)^35–37^. In contrast, a recent report describes neutralizing mAbs that bind NiV-F at membrane-proximal epitopes within DI and DII, expanding the observed sites of vulnerability on the HNV-F surface^39^.

**Figure 8.**
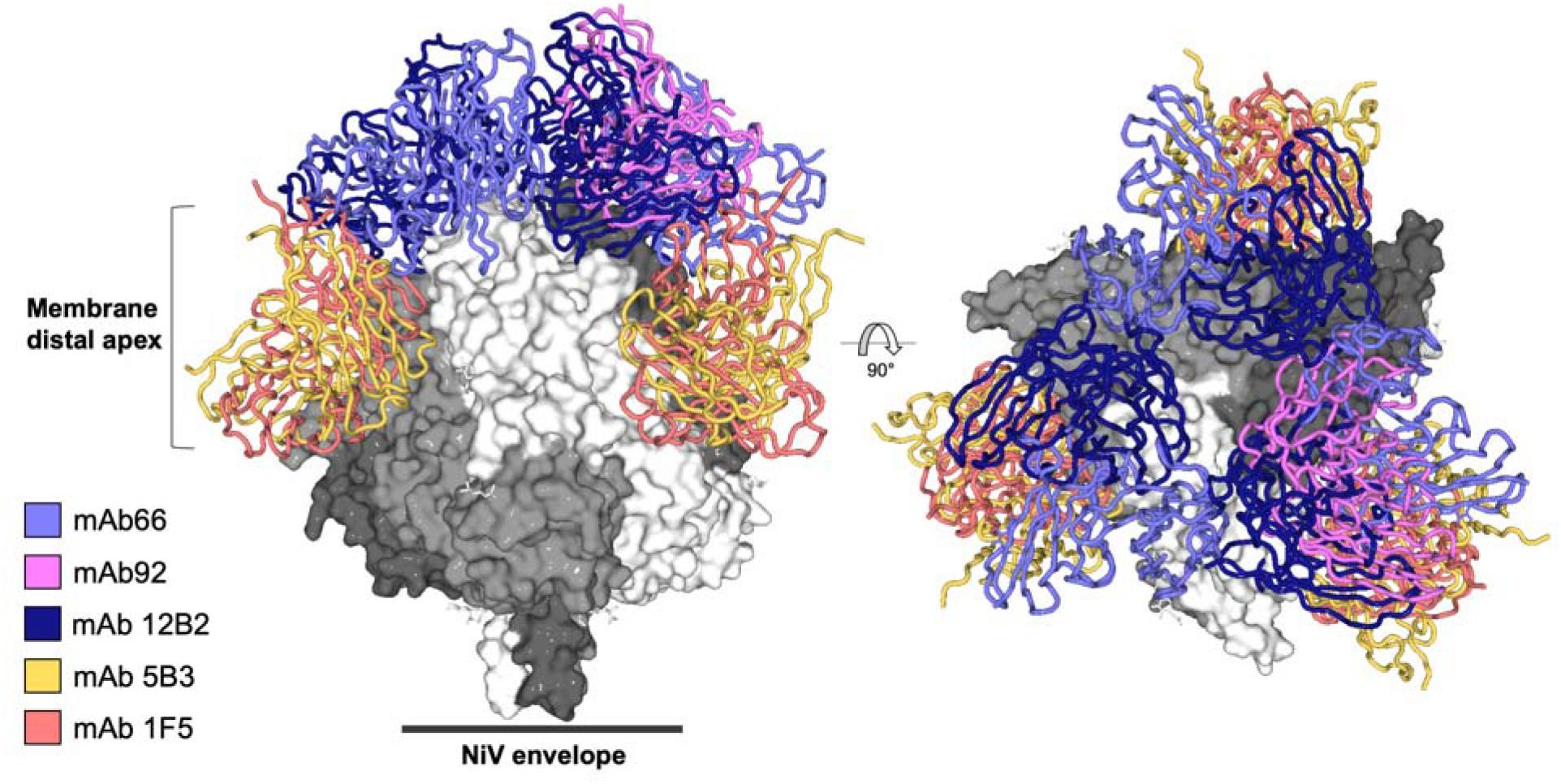
Mapping of membrane-distal mAb epitopes reveals the antigenic landscape of the HNV-F DII region. To visualize the breadth of characterized epitopes on NiV-F, the structures of mAbs in complex with HNV-F currently available in the PDB were modelled bound to the crystal structure of NiV-F (PDB 6T3F). The following structures are shown: mAb66 (light blue, PDB ID 6T3F^35^), mAb92 (light pink, PDB ID 7Z2F), mAb 5B3 (yellow, PDB ID 6TYS^36^), mAb 12B2 (dark blue, PDB ID 7KI4^37^), and mAb 1F5 (salmon, PDB 7KI6^37^). The NiV-F trimer is shown with surface representation, with each protomer colored a different shade of gray. The antibody fab molecules are depicted as cartoon tube. The relative position of the NiV-F viral envelop is noted with a black line.

The number of HNV mAbs and their epitopes thoroughly characterized in the literature is growing. Previously described structures of mAbs targeting HNV-G reveal epitopes including the dimerization interface (mAbs HENV-32 and HENV-103) and the receptor (ephrin) binding site (mAbs 102.3/102.4, HENV-26, and HENV-117)^31,32,38^. Antibodies targeting the membrane-distal DIII domain of HNV-F, mAbs 92, 66, 5B3, 15F, and 12B2^35–37^, have been shown to bind epitopes that include the HRA β-hairpin and helix (Fig 8) and are thought to act by stabilizing the fusion protein in its prefusion conformation, preventing membrane fusion from occurring. Though further experiments would be needed to fully elucidate the mechanism of neutralization by mAb92, it is plausible that mAb92 may also stabilize the prefusion state of the molecule, as the similar epitope of mAb66 has been shown to become disrupted in the postfusion state^35^. It is also worth noting that the F2 glycan (Asn67) is present within multiple apical epitopes and affects the binding of several of these mAbs to HNV-F, including mAb92. The recently described mAbs 4H3, 2D3, 1H8, 1F2, 1F3, and 4B8 also bind to the NiV-F apical DIII, highlighting the various neutralizing epitopes within this site of vulnerability. In contrast, mAbs 1H1, 1A9, 2B12, and 4F6 bind epitopes within DI and DII. Notably, the epitope of mAb 1A9 includes the fusion peptide, a novel epitope within paramyxoviruses^39^.

This initial study also showed that mAb92 offered complete protection against NiV disease when administered to Syrian hamsters prior to a lethal challenge with NiV Malaysia, demonstrating the mAb was effective *in vivo*. Syrian hamsters have been established as an effective and feasible model to study HNV disease, as they show similar respiratory and neurological signs to the disease that occurs in humans, making them an ideal model for the initial evaluation of therapeutics and vaccines^26,48–50^. Indeed, previous studies have shown that anti-G and anti-F mAbs have protected hamsters from NiV and HeV disease when administered prophylactically and post-infection, even using doses as low as 180 μg anti-F mAb and 1.12 μg anti-G mAb^25,26^. In addition to mAb92, the humanized anti-F monoclonal antibody mAb 5B3 was also shown to be protective in a ferret model against both NiV Bangladesh and HeV disease when administered post-challenge^30^. Notably, while mAb92 appears to be specific for NiV and did not confer protection against HeV, mAbs 5B3, 1F5, and 12B2 show activity against both NiV and HeV^30,36,37^. Additionally, the HNV-G targeting mAb 102.4 prevents NiV and HeV disease in an NHP model, has been used on a compassionate basis in humans, and has recently been shown to be safe and well tolerated in a Phase 1 clinical trial in Australia^28,29,33,34^, highlighting the clinical utility of these therapeutic approaches. Given the effectiveness of mAb92 and mAb 5B3 in animal models, it is plausible that mAbs targeting HNV-F will also show promise in clinical developments, particularly in antibody cocktails. While escape mutants from some of these mAbs have been observed in cell culture^36,38^, there has been no evidence of escape in the *in vivo* studies, likely from the high doses of antibody used. However, combining antibodies that target distinct epitopes, such as anti-F mAbs with anti-G mAbs, would reduce the possibility of escape mutants arising during treatment.

Both NiV and HeV continue to pose a sustained threat to human health, and newly discovered HNVs and HNV-like viruses occupy a wide geographical range^51^. Despite this threat, there is currently a paucity of approved treatments and vaccines against HNV disease licensed for human use, highlighting the need to define epitopes on the HNV surface and locate sites of vulnerability to allow the development of countermeasures. This study contributes to the growing body of knowledge of how immune system can neutralize NiV via antibody mediated targeting of the fusion glycoprotein and offers a structure-based rationale for the design of vaccines and therapeutics targeting HNV-F. Further evaluation of the *in vivo* and cross-neutralizing activity of anti-F mAbs targeting the apical DIII domain of NiV-F, both alone and in combination with mAbs targeting spatially distinct epitopes, would be a logical future direction in the development of treatments and preventative measures for HNV disease.

## Materials and Methods

### Hybridoma culture and mAb92 purification

Hybridomas secreting mAb92 were cultured in hybridoma-SFM (Gibco) at 37°C with 5% humidity. After ten days, the antibody-containing media was harvested, clarified by centrifugation at 4,000 x g for 20 minutes, then sterile filtered through a 0.22 μM vacuum filter unit. The mAb was purified using five milliliter HiTrap Protein G HP columns (GE Healthcare) equilibrated in 20 mM sodium phosphate buffer, pH 7, as per the manufacturer’s protocol. Eluted mAb92 was further purified by size exclusion chromatography (SEC) using a Superdex 200 Increase 10/300 GL column (GE Healthcare) equilibrated in phosphate buffered saline (PBS), pH 7.4.

### mAb92 cleavage to isolate Fab92

Purified mAb92 was subject to papain cleavage to isolate the Fab region. Concentrated mAb92 was diluted in 10 mM L-cysteine, 10 mM EDTA in phosphate buffered saline (PBS) (reaction buffer). Papain (Sigma) was activated by diluting at a 1:10 ratio (v/v) in 100 mM L-cysteine in PBS an incubated at room temperature for 15 minutes, then further diluted 1:100 (v/v) in reaction buffer. The diluted antibody and papain solutions were mixed 1:1 (v/v) and the cleavage reaction allowed to proceed for four hours at 37°C. The reaction was then stopped by the addition of 0.3 M iodoacetamide in a 1:10 ratio (v/v) to a final concentration of 30 mM.

The cleaved Fab was purified by a combination of Protein G purification and SEC. The reaction was flowed over one milliliter HiTrap Protein G HP columns (GE Healthcare) equilibrated in PBS, pH 7.4, to bind Fc or uncleaved mAb. The flowthrough containing the isolated Fab was concentrated and further purified by SEC using a Superdex 200 Increase 10/300 column equilibrated in 10 mM Tris pH 8, 150 mM NaCl.

### Sequencing and cloning of mAb 92 variable regions from hybridoma cell line

Hybridomas were cultured in medium E (Clonacell) at 37°C with 5% humidity. When cells achieved appropriate density, total RNA was extracted using the Qiagen RNeasy mini kit, per the manufacturer’s protocols. Purified RNA was eluted in 30 μL of RNase-free H_2_O. cDNA was generated using the Superscript IV First-Strand Synthesis System (Invitrogen) with 5 μL input RNA and random hexamers according to manufacturer’s protocol. The variable regions of heavy and κ chains were amplified by PCR using previously described rabbit primers and PCR conditions^52^. PCR products were gel purified and cloned into heavy and kappa expression plasmids^52,53^ adapted from the pFUSE-rIgG-Fc and pFUSE2-CLIg-rK1 ampicillin resistant vectors (InvivoGen) using the Gibson Assembly Master Mix (New England Biolabs) following the manufacturer’s protocol. Heavy and kappa variable region sequences were determined using Sanger sequencing (Genewiz). Antibody heavy and light chain plasmids were cotransfected at a 1:1 ratio (1 ug total DNA per 1×10^6^ cells) into HEK293F cells (ThermoFisher) using PEI Max 40K (linear polyethylenimine hydrochloride, Polysciences). Antibody supernatant was harvested 6 days following transfection and purified using protein G affinity chromatography following the manufacturers protocol (GE Healthcare). Specificity of the cloned mAb was determined by ELISA.

### Protein production (NiV-F)

The ectodomain of NiV-F (residues G26 – D482) was cloned into the pHLSEC vector (REF) containing the C-terminal GCNt trimerization motif (MKQIEDKIEEILSKIYHIENEIARIKKLIGE) in place of the transmembrane and cytosolic domains, as described previously^54^. The protein was expressed via transient transfection of adherent HEK 293T cells grown in roller bottles in the presence of kifunensine, an α-mannosidase I inhibitor^55^. Five days post transfection, the supernatant was harvested, clarified by centrifugation (4,000 x g for 30 minutes at 4°C), and diafiltrated against 10 mM Tris pH 8, 150 mM NaCl using an AKTA Flux system (GE Healthcare). The protein was further purified by Ni-NTA immobilized metal-affinity chromatography (IMAC) using five milliliter His-Trap HP columns (GE Healthcare), followed by size exclusion chromatography (SEC) using a Superose 6 Increase 10/300 column (GE Healthcare), equilibrated with 10 mM Tris pH 8, 150 mM NaCl. The purification was monitored by SDS-PAGE analysis.

### Cryo-electron microscopy sample preparation and data collection

To generate the NiV-F–Fab92 complex, the two proteins were mixed in a 3.3:1 molar ratio of Fab92 to NiV-F and incubated at room temperature for one hour. The complex was then purified and excess Fab92 removed by SEC using a Superose 6 Increase 10/300 column (GE Healthcare), equilibrated with 10 mM Tris pH 8, 150 mM NaCl. The purified NiV-F–Fab92 complex was concentrated to 0.6 μg/mL and 3.5 μL was applied onto freshly glow-discharged Quantifoil R 1.2/1.3 (Cu 200) grids. Excess liquid was removed by Vitrobot filter paper with a 3.5 second blot time at 4°C and 100% relative humidity before immediately plunge-freezing in a liquid ethane/propane mixture using a Vitrobot Mark IV (Thermo Fisher). Grids were stored at liquid nitrogen temperature until data collection.

Data were collected on a Titan Krios cryo-transmission electron microscope (Thermo Fisher) operating at 300 kV with a Gatan K3 detector using SerialEM in super-resolution mode at the Electron Biology Imaging Centre (eBIC) at Diamond Light Source (Didcot, UK). A total of 1,421 movies were collected with a super-resolution pixel size of 0.5425 Å.

### Cryo-electron microscopy data processing

Alignment and motion correction were performed using the Relion 3.1 implementation of motion correction^56^ with five-by-five patch-based alignment. Frames were binned by a factor of two for a final pixel size of 1.085 Å. Further processing took place using cryoSPARC v3.2.0^41^. The contrast transfer function was determined using Gctf-v1.06^57^. Images were manually inspected and those with poor fit discarded. Particles were first picked using unbiased blob picking, which were then subject to rounds of 2D classification. Classes representing top and side views were selected for use as templates for additional particle picking, the particles from which were again subject to 2D classification. A total of 394,502 particles were selected for *ab initio* reconstruction and heterogenous 3D refinement. The particles from the best class, totaling 178,307, were then refined further with non-uniform refinement^42^ to generate the final map that was then filtered for model building. Global resolution was determined by the gold standard FSC = 0.143 using cryoSPARC v3.2.0.

### Cryo-electron microscopy model building and refinement

The NiV-F and Fab portions of the crystal structure of the NiV-F–Fab66 complex (PDB ID 6T3F)^35^ were individually rigid body fit into the NiV-F–Fab92 electron microscopy density using Chimera. Sequence modification and real-space refinement were performed using COOT^58^. N-linked glycans were built and refined using the carbohydrates plug-in within COOT^59^. PHENIX real-space refinement was utilized for further refinement^60,61^, using secondary structure and Ramachandran geometry restraints. The model was validated by COOT, Molprobity, and PHENIX comprehensive validation^58,60–63^.

### Plasmid construction and cell-based binding assay

Codon-optimized NiV and HeV-F and -G glycoprotein contained an AU1 or HA tag, respectively. All mutants (F2mut, 70+74mut, 70mut, and 74mut) were generated by overlap PCR and sequence verified prior to use in experiments. To remove the F2 N-linked glycosylation site, the F2mut was generated by making a conservative asparagine to glutamine mutation, as previously described ^44^. The flow-cytometry based binding assay was performed as previously described^35^. Briefly, HEK293T cells, maintained in DMEM supplemented with 10% FBS, were transfected with the indicated construct using PEI (Transporter 5, Polysciences), then cells were collected two days post transfection with 10 mM EDTA. Cells were subsequently stained with a 1:1000 dilution of mAb92 of pAb2489/2490, then stained with an anti-Rb-674 secondary antibody prior to flow cytometry using the Guava easyCyte. As previously described, pAb2489 and pAb2490 displayed equivalent reactivity against NiV and HeV wt, F2, and 70+74 constructs^35^.

### Pseudotyped particle production and neutralization assay

Codon-optimized HNV -G, wt, or mutant F protein constructs were utilized to produce HNV pseudotyped particles (HNVpp) as previously described^35,44^. Briefly, HEK293T cells were transfected with the indicated G or F construct, infected with VSVΔG with a Renilla luciferase reporter gene (VSVΔG-RLuc), washed with DPBS, collected two days post infection, clarified by centrifugation, and purified by ultracentrifugation through a 20% sucrose cushion. Purified NiV or HeV pseudoparticles (NiVpp or HeVpp) were resuspended in DPBS and aliquoted prior to storage in -80°C. NiVpp and HeVpp were tittered on U-87 MG glioblastoma cells seeded in a 96 well plate to identify dilutions within the dynamic response range of the R. luciferase assay. A single dilution of the indicated HNVpp was incubated with an equal volume of serially diluted mAb92 for 30 min at room temperature. This mix was used to infect U87 cells for 24hrs prior to processing for luciferase activity as previously described^35^.

### Ethics statement

The animal study was approved by the Institutional Animal Care and Use Committee (IACUC) at Rocky Mountain Laboratories. The experiment took place following the guidelines outlined in the NIH Guide for the Care and Use of Laboratory Animals, the Welfare Act, United States Department of Agriculture and the United States Public Health Service Policy on Humane Care and Use of Laboratory Animals (Protocol #2019-33E), in an Association for Assessment and Accreditation of Laboratory Animal Care-approved facility by certified staff. Work with infectious Nipah and Hendra virus under biosafety level 4 conditions was approved by the Institutional Biosafety Committee (IBC) and samples were inactivated and removed from high containment according to IBC-approved standard operating procedures.

### Cells and virus

The Nipah Malaysia (GenBank AF212302) and Hendra virus (GenBank AF017149) isolates were obtained from the Special Pathogens Branch at the Centers for Disease Control and Prevention (CDC, Atlanta, USA), or Public Health Agency, Winnipeg, Canada. Henipavirus isolates were propagated on Vero E6 cells grown in Dulbecco’s Modified Eagle Medium (DMEM) supplemented with 2% fetal bovine serum (Gibco), 1 mM L-glutamine (Gibco), 50 U/mL penicillin and 50 μg/mL streptomycin (Gibco) at 37°C and 5% CO_2_ (virus isolation medium). Infectious titers of henipavirus virus stocks were determined by end-point titration, reported as log_10_ 50% tissue culture infective dose (TCID_50_/mL) in Vero E6 cells, using the method of Spearman and Karber. Vero E6 cells were maintained in DMEM supplemented with 10% fetal bovine serum (Gibco), 1 mM L-glutamine (Gibco), 50 U/mL penicillin and 50 μg/mL streptomycin (Gibco) at 37°C and 5% CO_2_.

### mAb92 *in vivo* study design

A total of 44 female Golden Syrian hamsters, aged 4 – 6 weeks, were purchased from Envigo. All animals were bled via orbital sinus puncture eight days before antibody administration (-8 days post inoculation (DPI)). At -2 DPI, 22 animals received 500 μg each of purified mAb92 via the intramuscular (I.M.) route and 22 animals received sterile saline. The animals that were treated with mAb92 were bled at -1 DPI via orbital sinus puncture. At 0 DPI, all animals were challenged with 100 LD_50_ of NiV Malaysia (6.8 x 10^3^ TCID_50_)^48^ or HeV (6.0 x 10^2^ TCID_50_)^49^ intraperitoneally (I.P.). Each study group consisted of 11 hamsters: 11 mAb92 treated plus 11 saline animals for each NiV Malaysia and HeV challenge. Of these, four animals were euthanized at 4 (HeV) or 5 (NiV) DPI and the remaining seven animals per group were followed for 28 days post challenge.

Animals were observed for clinical signs following infection and euthanized when pre-determined criteria were met as defined in the ASP, such as >20% weight loss or severe signs of disease including labored breathing and paralysis. Weight was recorded daily up to 14 DPI and combined nasal-oral swabs were collected starting on 0 DPI, then every other day until 13 DPI. Upon euthanasia, blood and tissues were collected for virology, serology, and histology as approved by IACUC.

### RNA extractions and quantitative PCR

Combined nasal-oral swabs were collected with prewetted swabs into 1 mL of DMEM supplemented with 50 U/mL penicillin and 50 μg/mL streptomycin (Gibco). RNA was extracted from the swab samples using the QIAmp Viral RNA Mini Kit (Qiagen) adapted for the QIAcube HT automated system. Brain and lung tissues harvested from animals at necropsy were homogenized in RLT buffer and RNA extracted using the RNeasy kit (Qiagen). The eluted RNA from tissues and swabs was analyzed by one-step qRT-PCR using 5 μL input RNA with the QuantiFast Probe PCR kit (Qiagen) and primer/probe sets targeting the NiV Malaysia and HeV N gene. RNA standards with known copy number were included with each assay, in duplicate.

### Titration assay

Tissues and swabs positive by qRT-PCR were analyzed for infectious virus titer by end-point titration. Vero E6 cells were seeded in 96 well plates at a density of 1.5 x 10^4^ cells per well in DMEM supplemented with 10% fetal bovine serum (Gibco), 1 mM L-glutamine (Gibco), 50 U/mL penicillin and 50 μg/mL streptomycin (Gibco) and incubated overnight at 37°C and 5% CO_2_. Weighed sections of tissues were homogenized in 1 mL virus isolation medium with a stainless-steel bead. Tissue homogenates and swab media were diluted ten-fold in virus isolation medium and used to inoculate the Vero E6 cells. The plates were incubated at 37°C and 5% CO_2_ for five days, after which wells were assessed for cytopathic effect (CPE). Wells that demonstrated CPE were counted and the infectious titer was determined by the method of Spearman and Karber using four replicates per sample.

### Enzyme-linked immunosorbent assay

Purified NiV-F protein was diluted to a concentration of 1 μg/mL in PBS. Maxisorp plates (Nunc) were coated with 50 μL (50 ng) of protein per well and incubated overnight at 4°C. Plates were then washed three times with PBS + 0.1% Tween-20 (PBST) and blocked with 100 μL casein in PBS blocking buffer (Thermo Fisher) for one hour at room temperature. Plates were again washed three times with PBST. Serum samples were serially diluted, in duplicate, in casein in PBS blocking buffer (Thermo Fisher) and 100 μL was added to the wells and allowed to incubate at room temperature for one hour. Wells were washed four times with PBST. Goat anti-hamster IgG (H+L) HRP conjugated secondary antibody (SeraCare 5220-0371) was diluted 1:3,000 in casein in PBS blocking buffer (Thermo Fisher), and 100 μL was added to the wells and allowed to incubate for one hour at room temperature. Wells were then washed five times with PBST. Plates were developed using the KPL TMB 2-component peroxidase substrate kit (SeraCare) and the reaction stopped with the KPL stop solution (SeraCare). Plates were read at 450 nm. Positive and negative controls were included on each plate. The threshold for positivity was calculated as the average plus three times the standard deviation of negative control wells. Reported serology titers are the reciprocal value of the highest dilution of serum at which signal was observed above the calculated threshold.

### Histology and in situ hybridization

Tissues were fixed in 10% Neutral Buffered Formalin, with two changes, for a minimum of seven days. Tissues were placed in cassettes and processed with a Sakura VIP-6 Tissue Tek, on a 12-hour automated schedule, using a graded series of ethanol, xylene, and PureAffin. Embedded tissues were sectioned at 5 μm and dried overnight at 42°C prior to staining.

Detection of NiV and HeV viral RNA was performed using the RNAscope VS Universal HRP assay (Advanced Cell Diagnostics Inc.) as previously described^64^, according to the manufacturer’s instructions. Briefly, tissue sections were deparaffinized and pretreated with heat and protease before hybridization with target-specific probes for NiV (Advanced Cell Diagnostics Inc. #439259) or HeV (Advanced Cell Diagnostics Inc #410719) and detected using the Discovery mRNA purple HRP detection kit (Roche Tissue Diagnostics #760-255) on the Ventana Discovery ULTRA staining platform (Roche Tissue Diagnostics). A board-certified veterinary anatomic pathologist blinded to the study groups evaluated all tissue slides.

### Statistical analysis

Statistical significance was calculated using GraphPad Prism v.9.1.0. The Log-rank (Mantel-Cox) test was used for survival. Multiple unpaired t-tests using the Holm-Sidak method were used to compare the virus titers from tissues. Significance is denoted as follows: ns = p > 0.05; * = p ≤ 0.05; ** = p ≤ 0.01; *** = p ≤ 0.001; **** = p ≤ 0.0001.

### Data Deposition

The sharpened and unsharpened cryo-EM maps and atomic model have been deposited in the EMDB and the Protein Data Bank with the accession codes EMD-14464 and PDB 7Z2F, respectively.

## Acknowledgements

The authors would like to acknowledge Stephanie N. Seifert, Rhys Pryce, Vojtech Prazak, Neeltje van Doremalen, and Anita Mora for their excellent technical assistance. Electron microscopy provision was provided through eBIC (EM20223-17) and the OPIC Electron Microscopy Facility, a UK Instruct-ERIC Centre, which was founded by a Wellcome Trust JIF award (060208/Z/00/Z) and is supported by a Wellcome equipment grant (093305/Z/10/Z). Computation used the Oxford Biomedical Research Computing (BMRC) facility, a joint development between the Wellcome Centre for Human Genetics and the Big Data Institute supported by Health Data Research UK and the NIHR Oxford Biomedical Research Centre. The research was supported by the Wellcome Trust Core Award Grant 203141/Z/16/Z with additional support from the NIHR Oxford BRC. In addition, the work was supported by the Division of Intramural research of the National Institute of Allergy and Infectious Diseases and through the Defense Advanced Research Planning Agency and Preventing Emerging Pathogenic Threats program Cooperative Agreement D18AC00031.The views expressed are those of the authors and not necessarily those of the NHS, the NIHR, the NIAID or the Department of Health. This work was funded by Medical Research Council MR/L009528/1 and MR/S007555/1 (to T.A.B.), and MR/N002091/1 and MR/V031635/1 (to T.A.B. and K.J.D.). T.A.B. and B.L. acknowledge funding from NIH Grant R01 AI123449 held with Alex Freiberg. The Wellcome Centre for Human Genetics is supported by Grant 203141/Z/16/Z.

## Author Contributions

Conceptualization: V.A.A, T.A.B, and V.J.M

Performed research: V.A.A, T.B. K.C.Y, K.Y.O, K.M.W, R.R, T.T, K.J.D.

Analyzed data: V.A.A, T.B., K.C.Y., K.Y.O, H.D., R.S., G.S., B.L, T.A.B, V.J.M

Writing – original draft: V.A.A., T.A.B, and V.J.M.

Writing – review and editing: V.A.A., K.Y.O., T.A.B, V.J.M.

Data Curation: V.A.A., K.Y.O., G.S., T.A.B., V.J.M.

Supervision: B.L., T.A.B., and V.J.M.

## References

1 Eaton, B. T., Broder, C. C., Middleton, D. & Wang, L. F. Hendra and Nipah viruses: different and dangerous. Nat Rev Microbiol 4, 23–35, doi:10.1038/nrmicro1323 (2006).

2 Hossain, M. J. et al. Clinical presentation of nipah virus infection in Bangladesh. Clin Infect Dis 46, 977–984, doi:10.1086/529147 (2008).

3 Luby, S. P. et al. Recurrent zoonotic transmission of Nipah virus into humans, Bangladesh, 2001-2007. Emerg Infect Dis 15, 1229–1235, doi:10.3201/eid1508.081237 (2009).

4 Luby, S. P. & Gurley, E. S. Epidemiology of henipavirus disease in humans. Curr Top Microbiol Immunol 359, 25–40, doi:10.1007/82_2012_207 (2012).

5 Hsu, V. P. et al. Nipah virus encephalitis reemergence, Bangladesh. Emerg Infect Dis 10, 2082–2087, doi:10.3201/eid1012.040701 (2004).

6 Arunkumar, G. et al. Outbreak Investigation of Nipah Virus Disease in Kerala, India, 2018. J Infect Dis 219, 1867–1878, doi:10.1093/infdis/jiy612 (2019).

7 WHO. 2018 Annual review of diseases prioritized under the Research and Development Blueprint. (2018).

8 Bowden, T. A., Crispin, M., Jones, E. Y. & Stuart, D. I. Shared paramyxoviral glycoprotein architecture is adapted for diverse attachment strategies. Biochem Soc Trans 38, 1349–1355, doi:10.1042/BST0381349 (2010).

9 Bowden, T. A. et al. Structural basis of Nipah and Hendra virus attachment to their cell-surface receptor ephrin-B2. Nat Struct Mol Biol 15, 567–572, doi:10.1038/nsmb.1435 (2008).

10 Yin, H. S., Wen, X., Paterson, R. G., Lamb, R. A. & Jardetzky, T. S. Structure of the parainfluenza virus 5 F protein in its metastable, prefusion conformation. Nature 439, 38–44, doi:10.1038/nature04322 (2006).

11 Lamb, R. A. & Jardetzky, T. S. Structural basis of viral invasion: lessons from paramyxovirus F. Curr Opin Struct Biol 17, 427–436, doi:10.1016/j.sbi.2007.08.016 (2007).

12 Wong, J. J., Paterson, R. G., Lamb, R. A. & Jardetzky, T. S. Structure and stabilization of the Hendra virus F glycoprotein in its prefusion form. Proc Natl Acad Sci U S A 113, 1056–1061, doi:10.1073/pnas.1523303113 (2016).

13 Xu, K. et al. Crystal Structure of the Pre-fusion Nipah Virus Fusion Glycoprotein Reveals a Novel Hexamer-of-Trimers Assembly. PLoS Pathog 11, e1005322, doi:10.1371/journal.ppat.1005322 (2015).

14 Zamora, J. L. R. et al. Third Helical Domain of the Nipah Virus Fusion Glycoprotein Modulates both Early and Late Steps in the Membrane Fusion Cascade. J Virol 94, doi:10.1128/JVI.00644-20 (2020).

15 Pager, C. T., Wurth, M. A. & Dutch, R. E. Subcellular localization and calcium and pH requirements for proteolytic processing of the Hendra virus fusion protein. J Virol 78, 9154–9163, doi:10.1128/JVI.78.17.9154-9163.2004 (2004).

16 Pager, C. T., Craft, W. W., Jr., Patch, J. & Dutch, R. E. A mature and fusogenic form of the Nipah virus fusion protein requires proteolytic processing by cathepsin L. Virology 346, 251–257, doi:10.1016/j.virol.2006.01.007 (2006).

17 Diederich, S., Thiel, L. & Maisner, A. Role of endocytosis and cathepsin-mediated activation in Nipah virus entry. Virology 375, 391–400, doi:10.1016/j.virol.2008.02.019 (2008).

18 Murin, C. D., Wilson, I. A. & Ward, A. B. Antibody responses to viral infections: a structural perspective across three different enveloped viruses. Nat Microbiol 4, 734–747, doi:10.1038/s41564-019-0392-y (2019).

19 Mire, C. E. et al. Single injection recombinant vesicular stomatitis virus vaccines protect ferrets against lethal Nipah virus disease. Virol J 10, 353, doi:10.1186/1743-422X-10-353 (2013).

20 Mire, C. E. et al. A recombinant Hendra virus G glycoprotein subunit vaccine protects nonhuman primates against Hendra virus challenge. J Virol 88, 4624–4631, doi:10.1128/JVI.00005-14 (2014).

21 DeBuysscher, B. L., Scott, D., Marzi, A., Prescott, J. & Feldmann, H. Single-dose live-attenuated Nipah virus vaccines confer complete protection by eliciting antibodies directed against surface glycoproteins. Vaccine 32, 2637–2644, doi:10.1016/j.vaccine.2014.02.087 (2014).

22 van Doremalen, N. et al. A single-dose ChAdOx1-vectored vaccine provides complete protection against Nipah Bangladesh and Malaysia in Syrian golden hamsters. PLoS Negl Trop Dis 13, e0007462, doi:10.1371/journal.pntd.0007462 (2019).

23 Geisbert, T. W. et al. A single dose investigational subunit vaccine for human use against Nipah virus and Hendra virus. NPJ Vaccines 6, 23, doi:10.1038/s41541-021-00284-w (2021).

24 Amaya, M. & Broder, C. C. Vaccines to Emerging Viruses: Nipah and Hendra. Annu Rev Virol 7, 447–473, doi:10.1146/annurev-virology-021920-113833 (2020).

25 Guillaume, V. et al. Antibody prophylaxis and therapy against Nipah virus infection in hamsters. J Virol 80, 1972–1978, doi:10.1128/JVI.80.4.1972-1978.2006 (2006).

26 Guillaume, V. et al. Acute Hendra virus infection: Analysis of the pathogenesis and passive antibody protection in the hamster model. Virology 387, 459–465, doi:10.1016/j.virol.2009.03.001 (2009).

27 Bossart, K. N. et al. A neutralizing human monoclonal antibody protects against lethal disease in a new ferret model of acute nipah virus infection. PLoS Pathog 5, e1000642, doi:10.1371/journal.ppat.1000642 (2009).

28 Bossart, K. N. et al. A neutralizing human monoclonal antibody protects african green monkeys from hendra virus challenge. Sci Transl Med 3, 105ra103, doi:10.1126/scitranslmed.3002901 (2011).

29 Geisbert, T. W. et al. Therapeutic treatment of Nipah virus infection in nonhuman primates with a neutralizing human monoclonal antibody. Sci Transl Med 6, 242ra282, doi:10.1126/scitranslmed.3008929 (2014).

30 Mire, C. E. et al. A Cross-Reactive Humanized Monoclonal Antibody Targeting Fusion Glycoprotein Function Protects Ferrets Against Lethal Nipah Virus and Hendra Virus Infection. J Infect Dis 221, S471–S479, doi:10.1093/infdis/jiz515 (2020).

31 Dong, J. et al. Potent Henipavirus Neutralization by Antibodies Recognizing Diverse Sites on Hendra and Nipah Virus Receptor Binding Protein. Cell 183, 1536–1550 e1517, doi:10.1016/j.cell.2020.11.023 (2020).

32 Doyle, M. P. et al. Cooperativity mediated by rationally selected combinations of human monoclonal antibodies targeting the henipavirus receptor binding protein. Cell Rep 36, 109628, doi:10.1016/j.celrep.2021.109628 (2021).

33 Playford, E. G. et al. Safety, tolerability, pharmacokinetics, and immunogenicity of a human monoclonal antibody targeting the G glycoprotein of henipaviruses in healthy adults: a first-in-human, randomised, controlled, phase 1 study. Lancet Infect Dis 20, 445–454, doi:10.1016/S1473-3099(19)30634-6 (2020).

34 Sazzad, H. M. S. A trial for post-exposure prophylaxis against henipaviruses. Lancet Infect Dis 20, 387–388, doi:10.1016/S1473-3099(19)30687-5 (2020).

35 Avanzato, V. A. et al. A structural basis for antibody-mediated neutralization of Nipah virus reveals a site of vulnerability at the fusion glycoprotein apex. Proc Natl Acad Sci U S A 116, 25057–25067, doi:10.1073/pnas.1912503116 (2019).

36 Dang, H. V. et al. An antibody against the F glycoprotein inhibits Nipah and Hendra virus infections. Nat Struct Mol Biol 26, 980–987, doi:10.1038/s41594-019-0308-9 (2019).

37 Dang, H. V. et al. Broadly neutralizing antibody cocktails targeting Nipah virus and Hendra virus fusion glycoproteins. Nat Struct Mol Biol 28, 426–434, doi:10.1038/s41594-021-00584-8 (2021).

38 Xu, K. et al. Crystal structure of the Hendra virus attachment G glycoprotein bound to a potent cross-reactive neutralizing human monoclonal antibody. PLoS Pathog 9, e1003684, doi:10.1371/journal.ppat.1003684 (2013).

39 Byrne, P. O. et al. Structural basis for antibody recognition of vulnerable epitopes on Nipah virus F protein. bioRxiv, 2022.2006.2013.495706, doi:10.1101/2022.06.13.495706 (2022).

40 Aguilar, H. C. et al. Polybasic KKR motif in the cytoplasmic tail of Nipah virus fusion protein modulates membrane fusion by inside-out signaling. J Virol 81, 4520–4532, doi:10.1128/JVI.02205-06 (2007).

41 Punjani, A., Rubinstein, J. L., Fleet, D. J. & Brubaker, M. A. cryoSPARC: algorithms for rapid unsupervised cryo-EM structure determination. Nat Methods 14, 290–296, doi:10.1038/nmeth.4169 (2017).

42 Punjani, A., Zhang, H. & Fleet, D. J. Non-uniform refinement: adaptive regularization improves single-particle cryo-EM reconstruction. Nat Methods 17, 1214–1221, doi:10.1038/s41592-020-00990-8 (2020).

43 Martin, A. C. & Thornton, J. M. Structural families in loops of homologous proteins: automatic classification, modelling and application to antibodies. J Mol Biol 263, 800–815, doi:10.1006/jmbi.1996.0617 (1996).

44 Aguilar, H. C. et al. N-glycans on Nipah virus fusion protein protect against neutralization but reduce membrane fusion and viral entry. J Virol 80, 4878–4889, doi:10.1128/JVI.80.10.4878-4889.2006 (2006).

45 Garner, O. B. et al. Endothelial galectin-1 binds to specific glycans on nipah virus fusion protein and inhibits maturation, mobility, and function to block syncytia formation. PLoS Pathog 6, e1000993, doi:10.1371/journal.ppat.1000993 (2010).

46 Krissinel, E. & Henrick, K. Inference of macromolecular assemblies from crystalline state. J Mol Biol 372, 774–797, doi:10.1016/j.jmb.2007.05.022 (2007).

47 WHO. Nipah virus disease - India, <https://www.who.int/emergencies/disease-outbreak-news/item/nipah-virus-disease---india> (2021).

48 DeBuysscher, B. L. et al. Comparison of the pathogenicity of Nipah virus isolates from Bangladesh and Malaysia in the Syrian hamster. PLoS Negl Trop Dis 7, e2024, doi:10.1371/journal.pntd.0002024 (2013).

49 Rockx, B. et al. Clinical outcome of henipavirus infection in hamsters is determined by the route and dose of infection. J Virol 85, 7658–7671, doi:10.1128/JVI.00473-11 (2011).

50 Wong, K. T. et al. A golden hamster model for human acute Nipah virus infection. Am J Pathol 163, 2127–2137, doi:10.1016/S0002-9440(10)63569-9 (2003).

51 Drexler, J. F. et al. Bats host major mammalian paramyxoviruses. Nat Commun 3, 796, doi:10.1038/ncomms1796 (2012).

52 Allen, E. R. et al. A Protective Monoclonal Antibody Targets a Site of Vulnerability on the Surface of Rift Valley Fever Virus. Cell Rep 25, 3750–3758 e3754, doi:10.1016/j.celrep.2018.12.001 (2018).

53 McCoy, L. E. et al. Holes in the Glycan Shield of the Native HIV Envelope Are a Target of Trimer-Elicited Neutralizing Antibodies. Cell Rep 16, 2327–2338, doi:10.1016/j.celrep.2016.07.074 (2016).

54 Chan, Y. P. et al. Biochemical, conformational, and immunogenic analysis of soluble trimeric forms of henipavirus fusion glycoproteins. J Virol 86, 11457–11471, doi:10.1128/JVI.01318-12 (2012).

55 Chang, V. T. et al. Glycoprotein structural genomics: solving the glycosylation problem. Structure 15, 267–273, doi:10.1016/j.str.2007.01.011 (2007).

56 Zivanov, J., Nakane, T. & Scheres, S. H. W. A Bayesian approach to beam-induced motion correction in cryo-EM single-particle analysis. IUCrJ 6, 5–17, doi:10.1107/S205225251801463X (2019).

57 Zhang, K. Gctf: Real-time CTF determination and correction. J Struct Biol 193, 1–12, doi:10.1016/j.jsb.2015.11.003 (2016).

58 Emsley, P., Lohkamp, B., Scott, W. G. & Cowtan, K. Features and development of Coot. Acta Crystallogr D Biol Crystallogr 66, 486–501, doi:10.1107/S0907444910007493 (2010).

59 Emsley, P. & Crispin, M. Structural analysis of glycoproteins: building N-linked glycans with Coot. Acta Crystallogr D Struct Biol 74, 256–263, doi:10.1107/S2059798318005119 (2018).

60 Liebschner, D. et al. Macromolecular structure determination using X-rays, neutrons and electrons: recent developments in Phenix. Acta Crystallogr D Struct Biol 75, 861–877, doi:10.1107/S2059798319011471 (2019).

61 Adams, P. D. et al. PHENIX: a comprehensive Python-based system for macromolecular structure solution. Acta Crystallogr D Biol Crystallogr 66, 213–221, doi:10.1107/S0907444909052925 (2010).

62 Davis, I. W. et al. MolProbity: all-atom contacts and structure validation for proteins and nucleic acids. Nucleic Acids Res 35, W375–383, doi:10.1093/nar/gkm216 (2007).

63 Chen, V. B. et al. MolProbity: all-atom structure validation for macromolecular crystallography. Acta Crystallogr D Biol Crystallogr 66, 12–21, doi:10.1107/S0907444909042073 (2010).

64 Wang, F. et al. RNAscope: a novel in situ RNA analysis platform for formalin-fixed, paraffin-embedded tissues. J Mol Diagn 14, 22–29, doi:10.1016/j.jmoldx.2011.08.002 (2012).

65 Crispin, M., Yu, X. & Bowden, T. A. Crystal structure of sialylated IgG Fc: implications for the mechanism of intravenous immunoglobulin therapy. Proc Natl Acad Sci U S A 110, E3544–3546, doi:10.1073/pnas.1310657110 (2013).

66 Corpet, F. Multiple sequence alignment with hierarchical clustering. Nucleic Acids Res 16, 10881–10890, doi:10.1093/nar/16.22.10881 (1988).

67 Robert, X. & Gouet, P. Deciphering key features in protein structures with the new ENDscript server. Nucleic Acids Res 42, W320–324, doi:10.1093/nar/gku316 (2014).

