## supplemental files for "A monoclonal antibody targeting the Nipah virus fusion glycoprotein apex imparts protection from disease"

**Supplementary Table S1.** Cryo-EM data collection and refinement statistics for NiV-F–Fab92.

|  | **NiV-F–Fab92** |
| --- | --- |
| **Data collection and processing** |  |
| Voltage (kV) | 300 |
| Electron exposure (e^-^/ Å^2^) | 44.37 |
| Defocus (μM) | 1.4 – 3.7 μM |
| Pixel size (Å/pix) | 0.5425 super-resolution |
| Symmetry | C1 |
| Final particle images (no.) | 178,307 |
| Map resolution (Å)^a^ | 3.5 |
| FSC threshold | 0.143 |
| **Refinement** |  |
| Initial model used | 6T3F |
| Model resolution (masked) (Å) | 3.0/3.3/3.5 |
| FSC threshold | 0/0.143/0.5 |
| Map sharpening B factor (Å^2^) | -155.35 |
| Model composition |  |
| Non-hydrogen atoms | 11924 |
| Protein residues | 1540 |
| Ligands | 12 |
| B factors (Å^2^) |  |
| Protein | 16.59/98.57/40.13 |
| Ligand | 41.90/69.21/53.05 |
| R.m.s deviations^b^ |  |
| Bond lengths (Å) | 0.004 |
| Bond angles (º) | 0.710 |
| **Validation** |  |
| MolProbity score^c^ | 1.50 |
| Clashscore | 4.81 |
| Poor rotamers (%) | 0.30 |
| Ramachandran^d^ |  |
| Favored (%) | 96.24 |
| Allowed (%) | 3.76 |
| Disallowed (%) | 0 |

^a^Gold standard FSC=0.143 from cryosparc v3.2.0^1,2^

^b^RMS deviations: root mean square deviation from ideal geometry.
^c/d^Ramachandran analysis determined with the Molprobity server^3,4^.

**Supplementary table S2.** Surviving hamster anti-NiV antibodies.

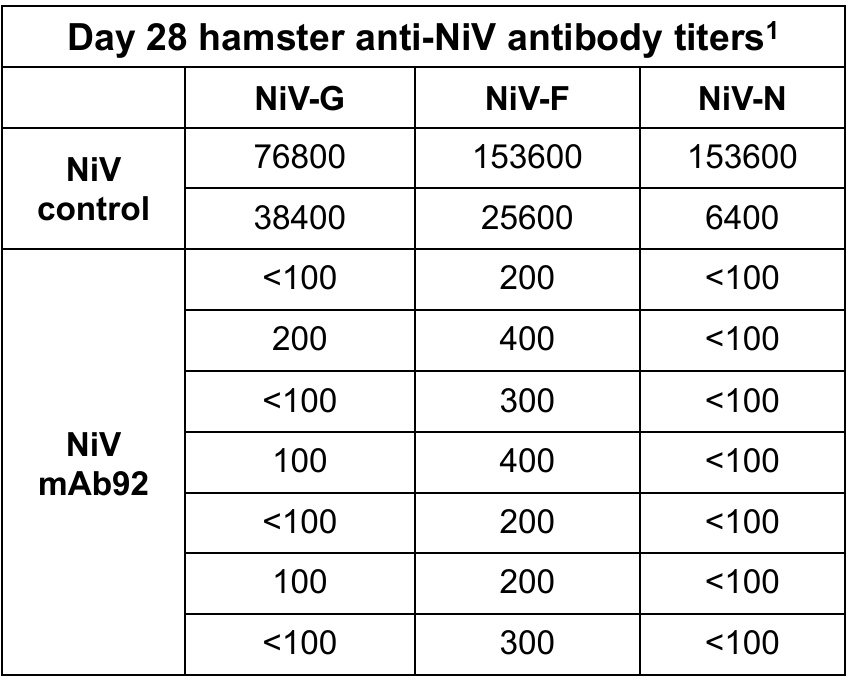

^1^Hamster IgG titers against NiV-G, F, and N proteins at 28 DPI determined by ELISA assay using an anti-hamster secondary antibody. Each sample was tested in duplicate and each row contains the end-point titers for an individual hamster.

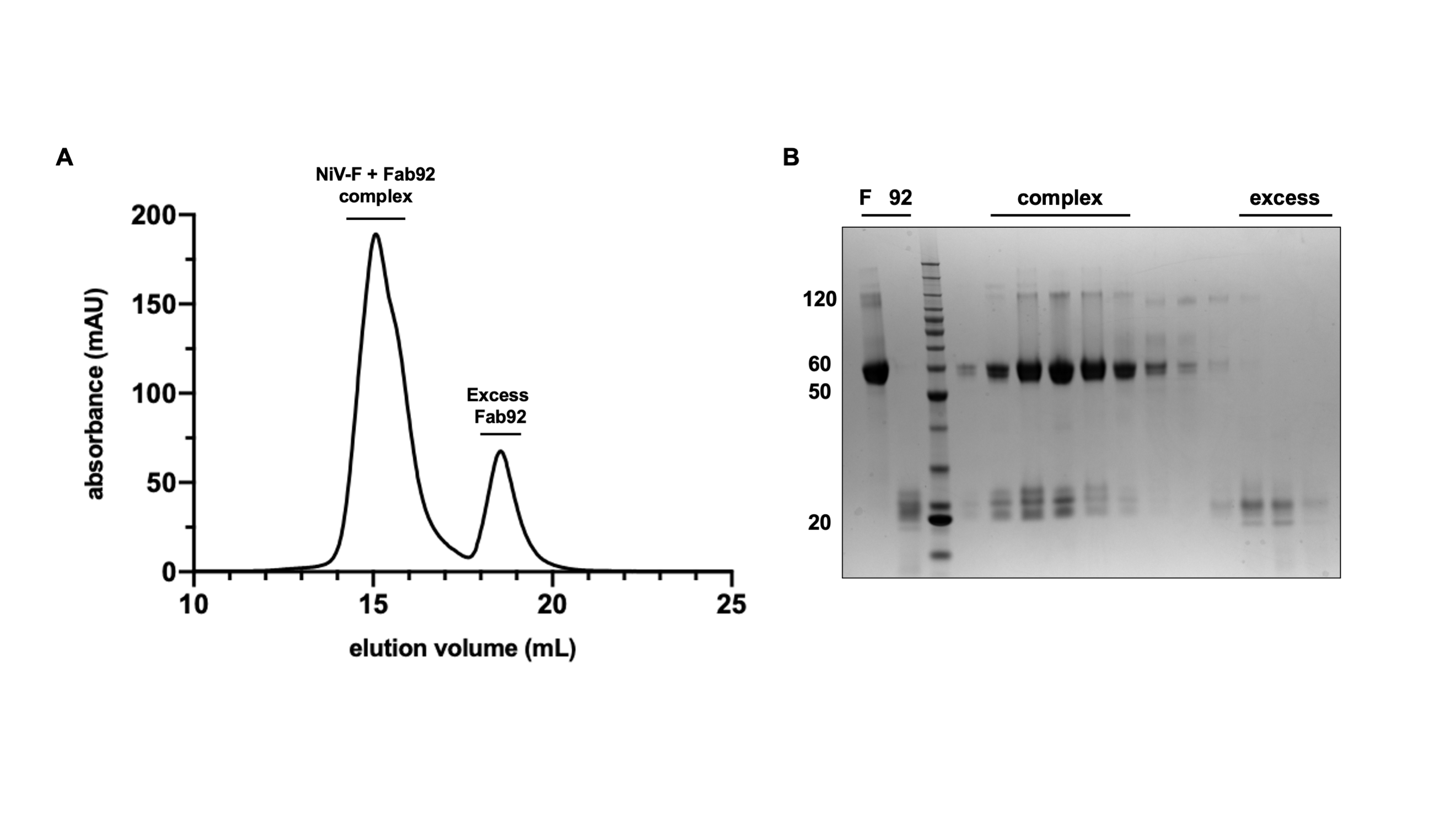

**Figure S1.** **NiV-F–Fab92 complex formation and purification.** (A) SEC chromatogram of NiV-F–Fab92 complex formation, with excess Fab92, using a Superose 6 Increase 10/300 column equilibrated in 10 mM Tris pH 8, 150 mM NaCl. (B) SDS-PAGE analysis of the eluted fraction of the NiV-F–Fab92 complex purification from panel (A) under reducing conditions. NiV-F and Fab92 are included individually to the left of the protein marker. Complex fractions (noted by a black line) were pooled for structural analysis.

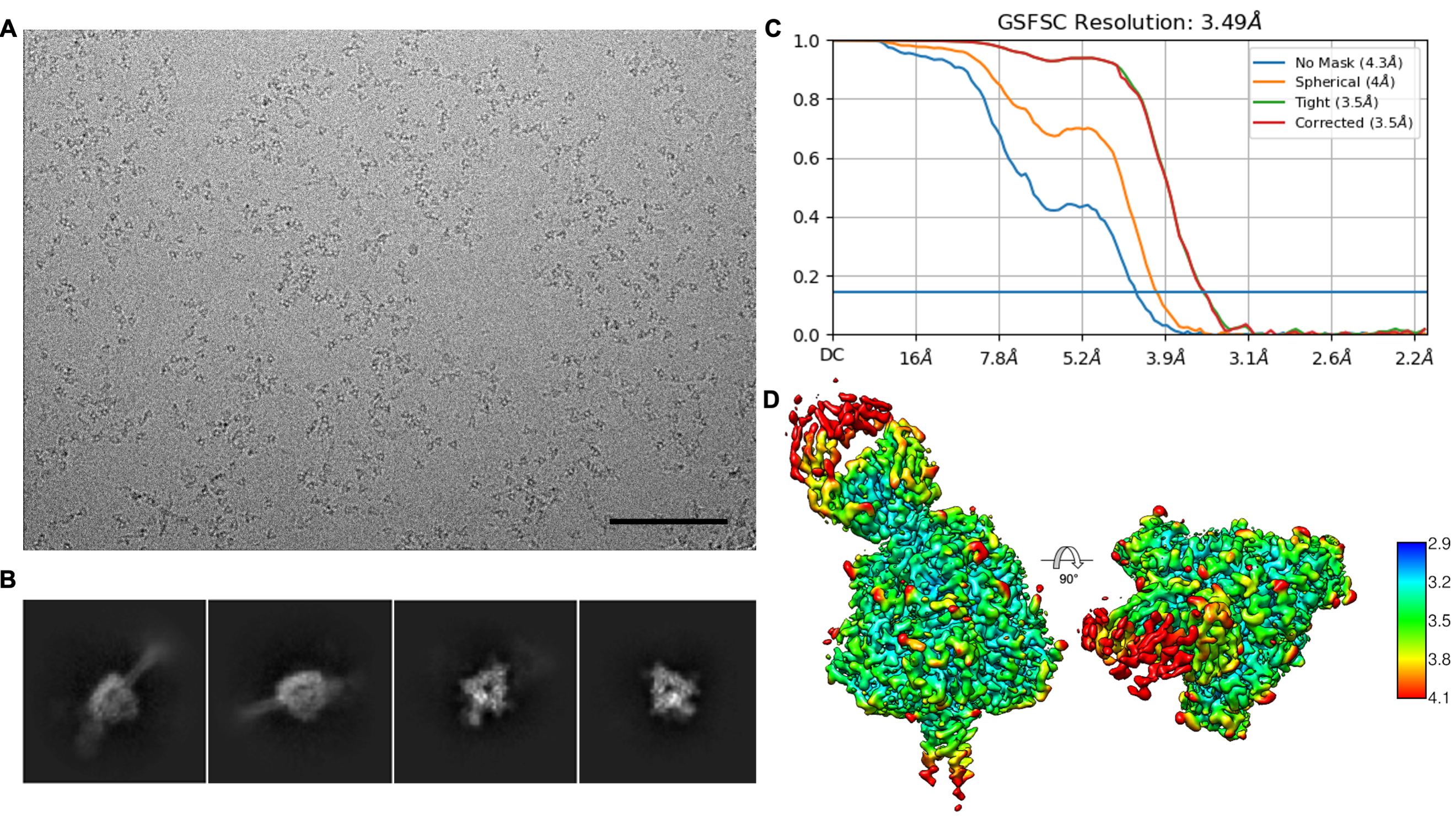

**Fig S2.** **Cryo-EM characterization of the NiV-F–Fab92 complex.** (A) Representative micrograph. Scale bar = 100 nm. (B) Representative selected 2D-class averages of side and top views. (C) Gold-standard Fourier shell correlation (FSC) plot, calculated by cryoSPARC v3.2.0^1,2^. The blue horizontal line represents the 0.143 threshold. (D) Side (left) and top (right) view of the NiV-F–Fab92 cryo-EM reconstruction colored by local resolution, using cryoSPARC v3.2.0 and Chimera. The colored scalebar represents local resolution determined by the cryoSPARC LocRes module, Å.

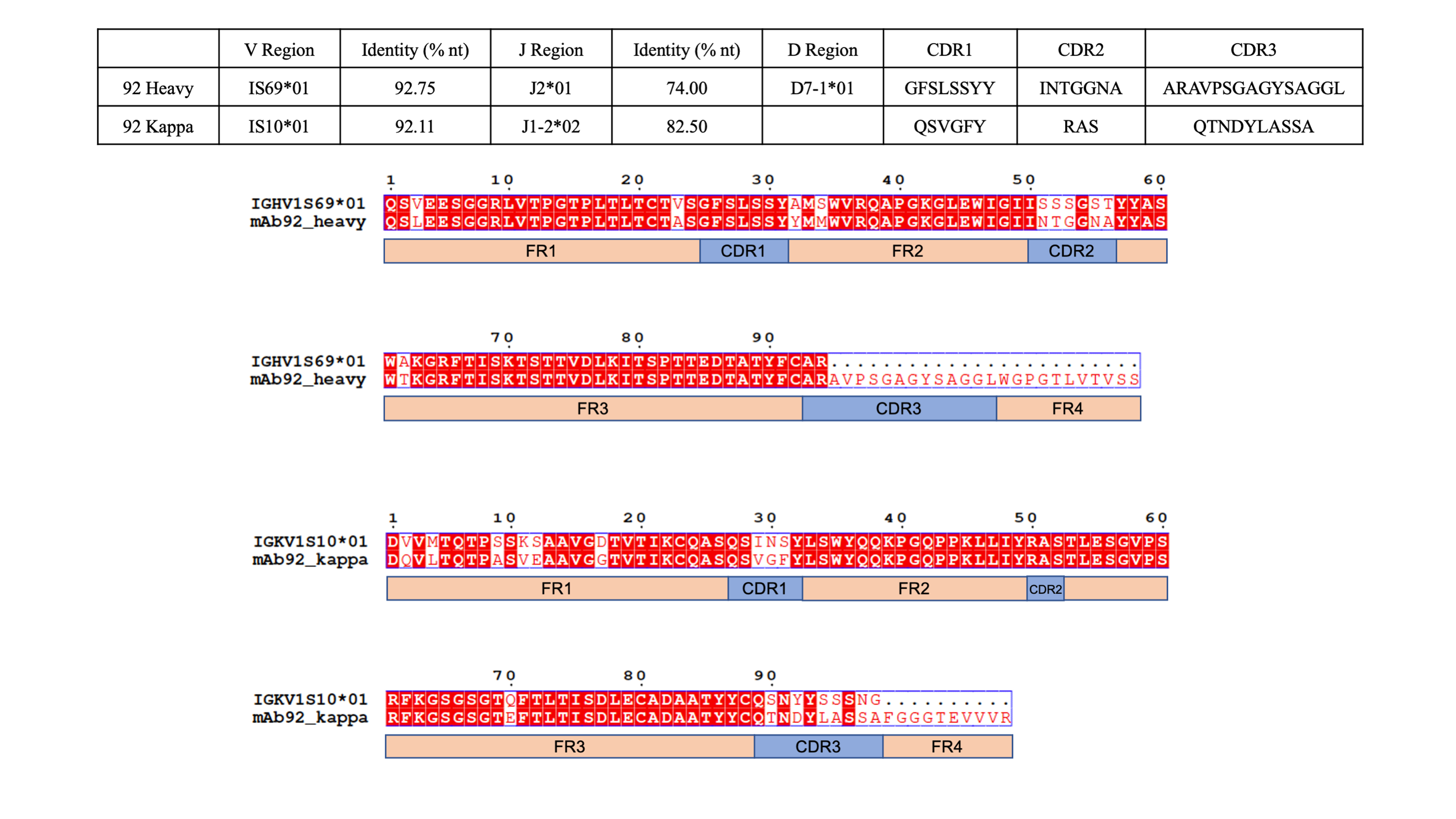

**Fig S3.** **Comparison of the rescued Fab92 sequence to the germline.** (Top) The international immunogenetics information system (IMGT) database^5,6^ was used to identify the most similar variable gene hits for both the heavy and kappa chain. (Bottom). The Fab92 heavy and kappa chain sequences were aligned using Multalin^7^ and displayed with ESPript^8^. The CDR and framework regions are shown by boxes under the alignment.

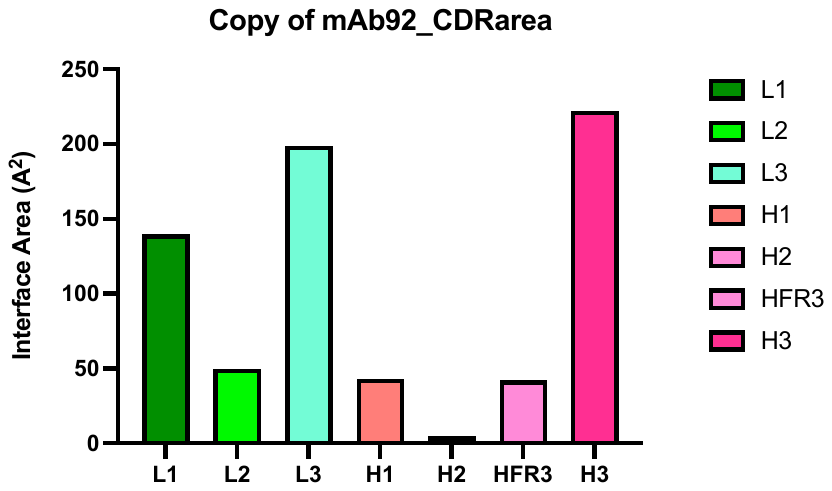

**Fig S4**. **Contributions of each Fab92 CDR loop to the proteinaceous interface.** The interface area was calculated by the PDBePISA server^9^, measured as interface in Å^2^.

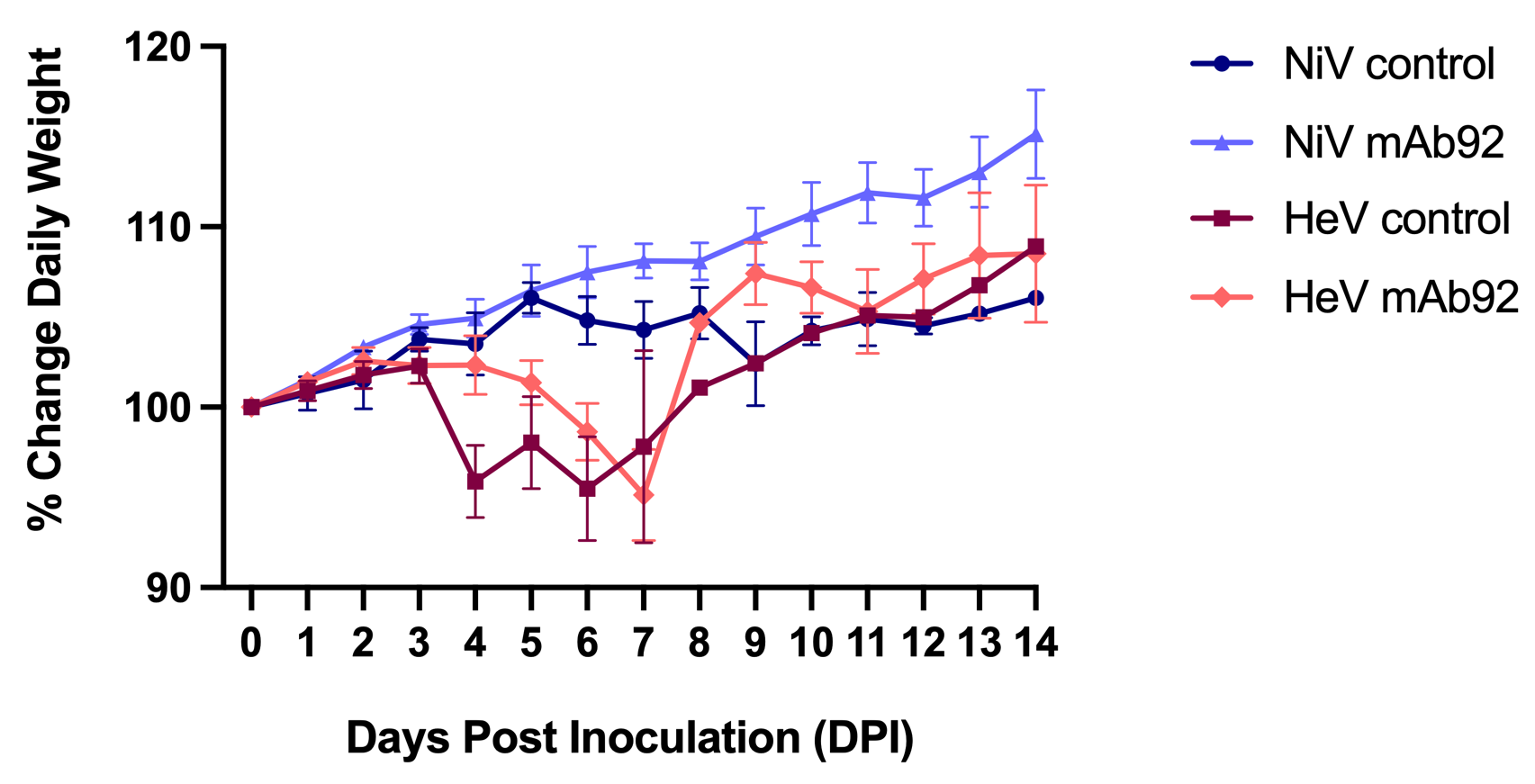

**Fig S5. Daily weights of hamsters following henipavirus challenge.** Control hamsters and mAb92 treated hamsters were weighed daily for 14 days following challenge. The average and standard error of the mean of the animals in each group is plotted. Each group consisted of seven animals, though group sizes decreased as animals were euthanized as endpoint criteria were met.

1	Punjani, A., Rubinstein, J. L., Fleet, D. J. & Brubaker, M. A. cryoSPARC: algorithms for rapid unsupervised cryo-EM structure determination. *Nat Methods* **14**, 290-296, doi:10.1038/nmeth.4169 (2017).

2	Punjani, A., Zhang, H. & Fleet, D. J. Non-uniform refinement: adaptive regularization improves single-particle cryo-EM reconstruction. *Nat Methods* **17**, 1214-1221, doi:10.1038/s41592-020-00990-8 (2020).

3	Davis, I. W. *et al.* MolProbity: all-atom contacts and structure validation for proteins and nucleic acids. *Nucleic Acids Res* **35**, W375-383, doi:10.1093/nar/gkm216 (2007).

4	Chen, V. B. *et al.* MolProbity: all-atom structure validation for macromolecular crystallography. *Acta Crystallogr D Biol Crystallogr* **66**, 12-21, doi:10.1107/S0907444909042073 (2010).

5	Brochet, X., Lefranc, M. P. & Giudicelli, V. IMGT/V-QUEST: the highly customized and integrated system for IG and TR standardized V-J and V-D-J sequence analysis. *Nucleic Acids Res* **36**, W503-508, doi:10.1093/nar/gkn316 (2008).

6	Giudicelli, V., Brochet, X. & Lefranc, M. P. IMGT/V-QUEST: IMGT standardized analysis of the immunoglobulin (IG) and T cell receptor (TR) nucleotide sequences. *Cold Spring Harb Protoc* **2011**, 695-715, doi:10.1101/pdb.prot5633 (2011).

7	Corpet, F. Multiple sequence alignment with hierarchical clustering. *Nucleic Acids Res* **16**, 10881-10890, doi:10.1093/nar/16.22.10881 (1988).

8	Robert, X. & Gouet, P. Deciphering key features in protein structures with the new ENDscript server. *Nucleic Acids Res* **42**, W320-324, doi:10.1093/nar/gku316 (2014).

9	Krissinel, E. & Henrick, K. Inference of macromolecular assemblies from crystalline state. *J Mol Biol* **372**, 774-797, doi:10.1016/j.jmb.2007.05.022 (2007).
